# Platelet PI3Kβ regulates breast cancer metastasis

**DOI:** 10.1101/2024.09.10.612261

**Authors:** Ryan C Graff, Adam Haimowitz, Jennifer T Aguilan, Adriana Levine, Jinghang Zhang, Wenlin Yuan, Merone Roose-Girma, Somasekar Seshagiri, Steven A Porcelli, Matthew J Gamble, Simone Sidoli, Anne R Bresnick, Jonathan M Backer

**Author notes:** Address correspondence to: Anne R Bresnick, Jonathan M Backer.

## Abstract

Platelets promote tumor metastasis by several mechanisms. Platelet-tumor cell interactions induce the release of platelet cytokines, chemokines, and other factors that promote tumor cell epithelial-mesenchymal transition and invasion, granulocyte recruitment to circulating tumor cells (CTCs), and adhesion of CTCs to the endothelium, assisting in their extravasation at metastatic sites. Previous studies have shown that platelet activation in the context of thrombus formation requires the Class IA PI 3-kinase PI3Kβ. We now define a role for platelet PI3Kβ in breast cancer metastasis. Platelet PI3Kβ is essential for platelet-stimulated tumor cell invasion through Matrigel. Consistent with this finding, *in vitro* platelet-tumor cell binding and tumor cell-stimulated platelet activation are reduced in platelets isolated from PI3Kβ mutant mice. RNAseq and proteomic analysis of human breast epithelial cells co-cultured with platelets revealed that platelet PI3Kβ regulates the expression of EMT and metastasis-associated genes in these cells. The EMT and metastasis-associated proteins PAI-1 and IL-8 were specifically downregulated in co-cultures with PI3Kβ mutant platelets. PI3Kβ mutant platelets are impaired in their ability to stimulate YAP and Smad2 signaling in tumor cells, two pathways regulating PAI-1 expression. Finally, we show that mice expressing mutant PI3Kβ show reduced spontaneous metastasis, and platelets isolated from these mice are less able to stimulate experimental metastasis in WT mice. Taken together, these data support a role for platelet PI3Kβ in promoting breast cancer metastasis and highlight platelet PI3Kβ as a potential therapeutic target.

**Significance:** We demonstrate that platelet PI3Kβ regulates metastasis, broadening the potential use of PI3Kβ-selective inhibitors as novel agents to treat metastasis.

## Introduction

Metastasis is the primary cause of mortality in patients with solid tumors. Platelets promote metastasis via direct interactions with circulating tumor cells (CTCs) [1–5]. Tumor cell binding activates platelets, leading to the release of platelet granule contents that promote tumor cell invasion and metastasis [6, 7]. These include TGF-β, which promotes epithelial-mesenchymal transition (EMT) in tumor cells [6, 8]. Inhibition of platelet activation or platelet TGF-β release prevents platelet-mediated tumor cell EMT and metastasis [6, 9].

Platelet activation, adhesion, and *in vivo* thrombus formation require the activity of the Class IA PI 3-kinase PI3Kβ [10–14]. Platelet activation by collagen (glycoprotein VI and integrin α2β1), thrombin (PAR1), and ADP (P2Y12) is disrupted by inhibition or knockout of platelet PI3Kβ [11, 13, 15, 16]. PI3Kβ produces the majority of PIP_3_ in platelets upon stimulation by thrombin and collagen-related peptide (CRP), suggesting that PI3Kβ plays a unique role in classical platelet activation [17]. More recent studies have shown that platelet PI3Kβ is essential for mediating platelet-leukocyte interactions in the context of bacterial pneumonia, suggesting a role for PI3Kβ in mediating the platelet response to disease [18].

Activation of PI3Kβ requires simultaneous inputs from receptor tyrosine kinases and G-protein coupled receptors (GPCRs) [19]. Our lab previously identified the Gβγ binding site in p110β [20]. Mutation of this site (p110β^526/527^KKDD, referred to as PI3Kβ^KKDD^) selectively disrupts GPCR coupling to PI3Kβ and abolishes PI3Kβ activation in myeloid cells [20, 21]. Importantly, PI3Kβ^KKDD^ still exhibits basal lipid kinase activity, and the PI3Kβ^KKDD^ mutation is not equivalent to a knock-out or kinase-deficient mutant. However, expression of PI3Kβ^KKDD^ in breast cancer cells inhibits matrix degradation, invasion, macropinocytosis, and experimental metastasis [22–24].

Although the role of PI3Kβ in the hemostatic function of platelets is well described, its role in platelet-mediated cancer metastasis is unexplored. Recent studies examining platelet-tumor cell interactions suggest potential roles for PI3Kβ. Platelet integrin α2β1, integrin α6β1, glycoprotein VI, and CLEC-2 are all implicated in platelet-tumor cell interactions [2, 4, 5, 25], and their activation of platelets via ITAM signaling is dependent on PI3Kβ [4, 11]. Platelet PI3Kβ also mediates inside-out activation of integrins, potentially enhancing platelet-tumor cell interactions [11, 14, 16, 26].

In this study, we demonstrate a requirement for platelet PI3Kβ in breast cancer metastasis. Mutation of platelet PI3Kβ reduces platelet activation in response to tumor cell binding. PI3Kβ^KKDD^ platelets are impaired with regard to the induction of tumor cell EMT, invasiveness, and metastasis. These data suggest that platelet PI3Kβ is a novel target in metastasis, potentially broadening the application of PI3Kβ-selective inhibitors in cancer chemotherapy.

## Methods

### Generation of PI3KβKKDD mice

Mice carrying the *PIK3CB^526/527^KKDD* allele were generated using C57BL/6N ES cells as previously described [27]. Fertilized, one-celled embryos (zygotes) were microinjected on E0.5. The *PIK3CB^526/527^KKDD* allele was produced by inserting a cassette containing a *LoxP*, wild-type *PIK3CB* cDNA (exons 13-24), a human growth hormone 3′UTR followed by a 4x polyadenylation (polyA) signal, an *FRT-Neo^R^-FRT* selection marker and a second *LoxP*, into the *PIK3CB* locus 5′ of a mutated exon 13 encoding the KK^526/527^DD (AAAAAA to GACGAC) change (Figure S1). The *Neo^R^* cassette was excised in ES cells using adeno-FLP prior to microinjection. *PIK3CB^526/527^KKDD* mice were maintained on a C57BL/6N genetic background.

### Platelet isolation and tumor cell co-culture

Platelets were isolated from 8–20-week-old mice. Blood was collected via cardiac puncture into a syringe containing acid citrate dextrose buffer (85 mM trisodium citrate, 71 mM citric acid, 111 mM glucose, pH 4.5), and decanted into a fresh tube containing Modified Tyrode-HEPES buffer (MTHB) (138 mM NaCl, 10 mM HEPES, 12 mM sodium bicarbonate, 2.7 mM KCl,

0.4 mM sodium phosphate monobasic, 5 mM glucose, 0.35% (w/v) BSA, pH 7.4). The samples were centrifuged at 200xg for 15 min and allowed deaccelerate with no brake. The supernatant (platelet-rich plasma) was transferred to a fresh tube. PGI_2_ (Millipore Sigma; P6188) and apyrase (Millipore Sigma; A6535) were added to final concentrations of 1 µg/mL and 0.02 U/mL, respectively. Platelets were pelleted by centrifugation at 1000xg for 5 min, resuspended in MTHB supplemented with 0.02 U/mL apyrase, and counted (Genesis hematology analyzer, Oxford Sciences). For tumor cell co-cultures, platelets were added to tumor cells at a ratio of 50 platelets:1 tumor cell and incubated for 12-40 hours depending on the experiment.

### Human platelet isolation

Human blood collection was performed with Institutional Review Board approval from Albert Einstein College of Medicine (082099). Volunteers did not take platelet inhibitors for at least two weeks prior to blood collection. Donors were all healthy males between 25-65 years old. Blood was collected from the median cubital vein by standard venipuncture methods into syringes containing acid citrate dextrose buffer. Platelets were then isolated and counted as described above.

### Experimental metastasis

GFP-E0771 or Cerulean-PyMT cells were co-cultured without or with platelets isolated from WT or PI3Kβ^KKDD^ mice as described above. After 40 hours, the cells were washed three times with PBS to remove platelets, detached with trypsin and washed twice more with PBS.

2.5x10^5^ tumor cells were injected into the tail vein of 10–16-week-old female C57Bl/6 mice. After 14 days, mice were euthanized with isoflurane and lungs were collected. The anterior and posterior aspects of the lungs from mice injected with GFP-E0771 cells were imaged using a Leica MZ FLIII stereoscope with a 1x objective. Surface metastases were quantitated with ImageJ (National Institutes of Health); the area of metastasis was measured by thresholding and expressed as a percent of the total lung surface area. Lungs were then minced and resuspended in TRIzol (Invitrogen; 15596026), and RNA was extracted from lung samples according to standard techniques. Reverse transcription of RNA samples was performed with SuperScript™ IV VILO™ Master Mix with ezDNase™ Enzyme (Invitrogen; 11766050). Assessment of lung metastatic burden was performed by quantitative PCR (qPCR) using PowerTrack™ SYBR Green Master Mix (Applied Biosystems; A46109). Samples were run on a QuantStudio™ 6 Flex Real-Time PCR System (Applied Biosystems; 4485691) with the following primers:

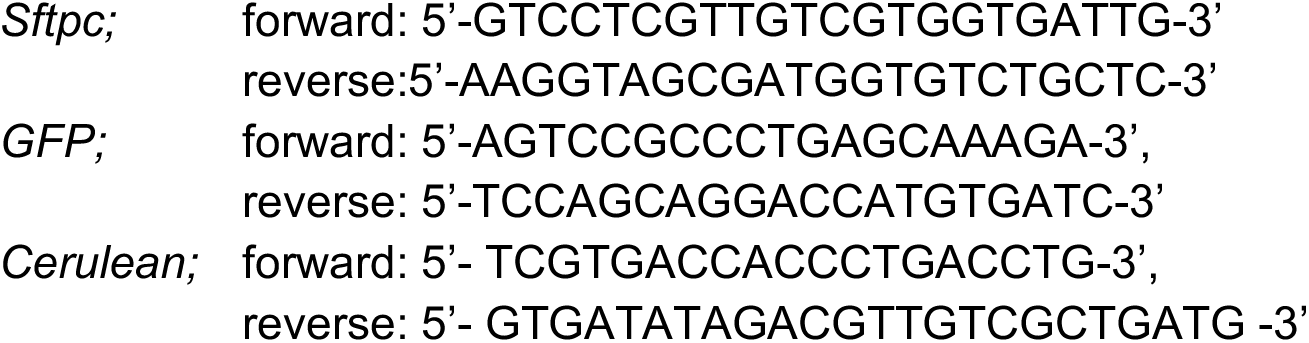

### Spontaneous metastasis

GFP-E0771 cells were detached with trypsin, washed three times with PBS, and resuspended in 50% PBS, 50% growth factor-reduced Matrigel (Corning; CB-40230A). 1.0x10^5^ cells were injected into the mammary fat pad of 12–16-week-old female WT or PI3Kβ^KKDD^ mice. Tumor size was monitored three times a week using digital calipers. Mice were euthanized when tumors reached a size of 1.5 cm^3^, or when body condition scores were ≤ 2. Blood was collected and analyzed by flow cytometry as described below. Lungs were isolated, minced, and resuspended in TRIzol. Lung metastatic burden was quantified by qPCR as described above.

### Flow cytometry analysis of blood from tumor-bearing mice

Blood was collected from GFP-E0771 orthotopic tumor-bearing mice via cardiac puncture into a syringe containing 500 mM EDTA, decanted into a fresh tube and analyzed with a hematology analyzer (Genesis, Oxford Sciences). RBC lysis buffer (0.02 M Tris-HCl, pH 7.2, 0.14 M NH_4_Cl) was added to whole blood for 10 min. Cells were then washed twice and resuspended in PBS with 4% FBS. Fc receptors were blocked using TruStain FcX (anti-mouse CD16/32) antibody (93) prior to surface staining with anti-mouse mAb against: CD41-AF647 (MWReg30), CD62P/P-selectin-PE-Cy7 (RMP-1), CD11b-BUV496 (M1/70), CD115-BV605 (AFS98), Ly-6G-PerCP(1A8), CD3-AF700 (17A2), CD45-APC-efluor780 (30-F11) at 4°C for 30 min. Single stain controls were performed using BD Compensation Particles Sets (BD Biosciences; 552845 and 552843). Cells were analyzed on an Aurora flow cytometer (Cytek Biosciences). Data was analyzed using FlowJo software (BD Biosciences).

### In vitro platelet activation and tumor cell binding

For ligand-stimulated platelet activation, platelets were isolated from WT or PI3Kβ^KKDD^ mice as described above and resuspended in MTHB supplemented with 2 mM calcium. Platelets were stimulated with collagen I (20 µg/mL; Corning; 354249) and ADP (20 µM; Fisher Scientific; A06261G) for 30 min at 37°C while rotating at 8 rpm. To measure tumor cell-stimulated platelet activation and tumor cell-platelet binding, tumor cells were labeled with CellTracker™ Green CMFDA Dye (Invitrogen; C7025) for 25 min at 37°C, pelleted, and resuspended in MTHB supplemented with 10% freshly prepared mouse serum. Tumor cells were then incubated without or with platelets from WT or PI3Kβ^KKDD^ mice at a 1:50 ratio for 30 min at 37°C with rotation as above. To measure binding and activation, Fc receptors were blocked as described above and cells were stained with anti-mouse mAb against CD41-PE (MWReg30) and CD62P/P-selectin- APC (Psel.KO2.3) at 4°C for 30 min. Samples were analyzed on an Aurora flow cytometer (Cytek Biosciences). Data was analyzed using FlowJo software (BD Biosciences).

### RNA sequencing

MCF10A cells were cultured for 40 hours without or with platelets isolated from WT or PI3Kβ^KKDD^ mice as described above. Cells were washed three times to remove platelets before lysis in TRIzol. Platelet removal was confirmed by FACS analysis of trypsinized cells from parallel dishes. RNA was extracted and evaluated for quality using the 5200 Fragment Analyzer system (Agilent). Only samples with an RNA Quality Number of 10.0 were sequenced. Sequencing was performed by MedGenome (40 million paired-end reads). Reads were aligned to the hg19 genome annotation using STAR [28], and featureCounts [29] was used to generate count tables as inputs for DEseq2, which was used to normalize the data [30]. The normalized data were analyzed by Gene Set Enrichment Analysis (GSEA) using the Molecular Signatures Database collection of Hallmark Gene Sets.

### Statistics

Data are expressed as the mean ± SEM from at least three independent experiments. All statistical analyses were performed using GraphPad Prism software. If the data was normally distributed, it was analyzed with a student’s t-test or one-way ANOVA. If the data was not normally distributed, it was analyzed using a Mann–Whitney *U* test or Kruskal–Wallis *H* test. Statistical analyses were designed in consultation with the Einstein Biostatistics Shared Resource.

Additional details are provided in Supplemental Methods.

## Results

### Platelet PI3Kβ signaling is required for metastasis

To investigate the contribution of stromal PI3Kβ signaling in breast cancer metastasis, we developed mice with a whole-body knock-in of the p110β^526/527^KKDD mutation (Figure S1). This mutant PI3Kβ (henceforth referred to as PI3Kβ^KKDD^) is defective for binding to Gβγ [20]. Given that activation of PI3Kβ requires coincident signaling from both RTKs and GPCRs [19, 21], the PI3Kβ^KKDD^ mutant has normal basal activity but is minimally activated in response to growth factors and/or cytokines.

GFP-E0771 murine mammary carcinoma cells were injected into the mammary fat pads of syngeneic WT or PI3Kβ^KKDD^ mice. Tumor sizes were monitored by caliper measurement and no differences in primary tumor growth were observed (Figure 1A). When mammary tumors reached 1.5 cm^3^, lungs were isolated and analyzed by qPCR for expression of GFP mRNA as a measure of metastatic burden. In both the left and right lungs, there was a clear trend toward a reduction in spontaneous metastasis in the PI3Kβ^KKDD^ mice (Figure 1B), suggesting an important role for stromal PI3Kβ in breast cancer metastasis.

**Figure 1.**
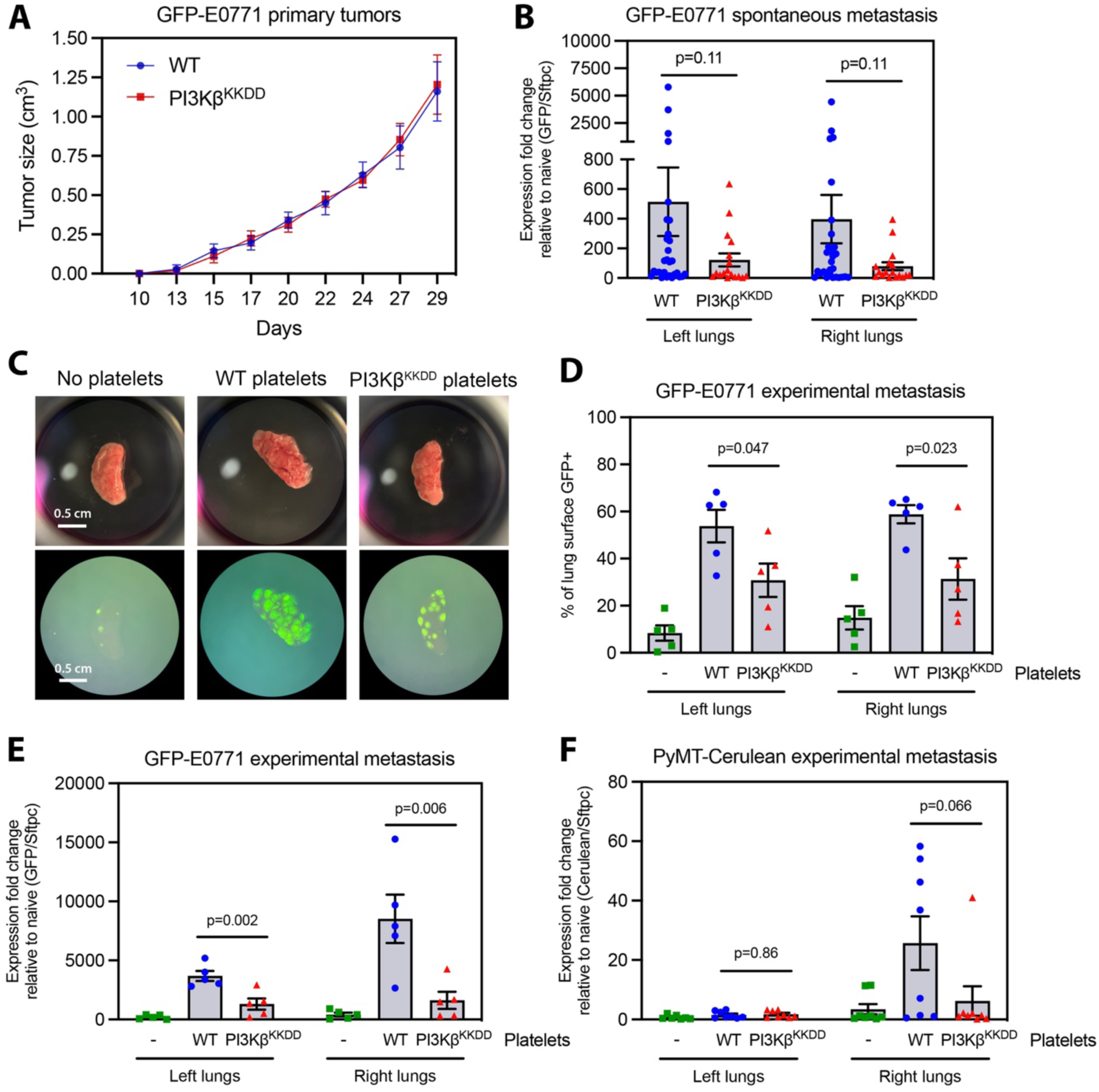
Platelet PI3Kβ activity is required for metastasis to lungs. (A) Growth of orthotopic syngeneic GFP-E0771 tumors in WT or PI3Kβ^KKDD^ C57Bl/6 mice. (B) qPCR quantitation of GFP expression in the left and right lungs. Data are presented as the mean ± SEM from 9-10 (A) or 17-29 (B) mice per group. (C) GFP-E0771 tumor cells were co-cultured with WT or PI3Kβ^KKDD^ platelets and injected into the tail veins of WT mice. Representative fluorescent images of left lungs removed after 14 days are shown. (D) Quantitation of the GFP+ lung surface area. (E) Quantitation of GFP expression in whole lungs by qPCR. Data are presented as the mean ± SEM from 5 mice per group. (F) Tumor cells expressing PyMT and Cerulean were co-cultured with WT or PI3Kβ^KKDD^ platelets and injected into the tail veins of WT mice and analyzed at 14 days as in (E). Data are presented as the mean ± SEM from 7-8 mice per group.

Given the established roles of platelets in metastasis [31] and of PI3Kβ activity in platelet activation [11–15, 17], we investigated the ability of PI3Kβ^KKDD^ platelets to promote metastasis of mammary tumor cells. Consistent with previous studies of PI3Kβ signaling during platelet activation in response to canonical platelet ligands [11–15, 17], PI3Kβ^KKDD^ platelets show reductions in spreading on fibrinogen (Figures S2A and S2B) and activation in response to ADP and soluble collagen I (Figures S2C and S2D). We next added WT or PI3Kβ^KKDD^ platelets to *in vitro* cultures of GFP-E0771 cells for 40 hrs, an established technique for inducing tumor cell epithelial-mesenchymal transition (EMT) and increasing the metastatic behavior of tumor cells [6]. After removal of the platelets, the GFP-E0771 cells were injected into the tail veins of WT mice and lungs were harvested 14 days later. Quantification of lung surface metastases by GFP fluorescent area, and quantification of GFP mRNA expression in whole lungs, revealed that GFP- E0771 cells cultured with PI3Kβ^KKDD^ platelets were less able to metastasize as compared to cells cultured with WT platelets (Figures 1 C-E). Similar results were obtained using Cerulean-labeled tumor cells from MMTV-PyMT transgenic mice cultured without or with WT or PI3Kβ^KKDD^ platelets (Figure 1F). In this case, tumor cells exclusively metastasized to the right lung for unknown reasons. Nonetheless, cells cultured with PI3Kβ^KKDD^ platelets show a clear trend toward reduced metastatic burden.

### Platelet-tumor cell interactions and tumor cell-stimulated platelet activation require platelet PI3Kβ

CTCs bind to platelets in the blood stream, resulting in platelet activation [6, 9, 32, 33]. To test the role of platelet PI3Kβ in platelet-tumor cell interactions, we incubated platelets and Cell Tracker-labeled tumor cells in suspension at different ratios and analyzed their interactions by flow cytometry (Figures 2A-D). PI3Kβ^KKDD^ platelets showed reduced binding to both human MDA-MB-231 breast adenocarcinoma cells and murine E0771 cells at low platelet-tumor cell ratios (Figures 2B and 2D), suggesting that mutation of PI3Kβ causes a right shift in the platelet-tumor cell binding curve. We also measured the activation of tumor cell-bound platelets by staining for cell surface P-selectin. We observed an approximately 2-fold reduction in the activation of tumor cell-bound PI3Kβ^KKDD^ platelets as compared to WT platelets (Figures 2E and 2F). In contrast, there was minimal activation of platelets from either genotype when not bound to tumor cells. These data show that platelet activation in response to direct tumor cell binding depends on platelet PI3Kβ.

**Figure 2.**
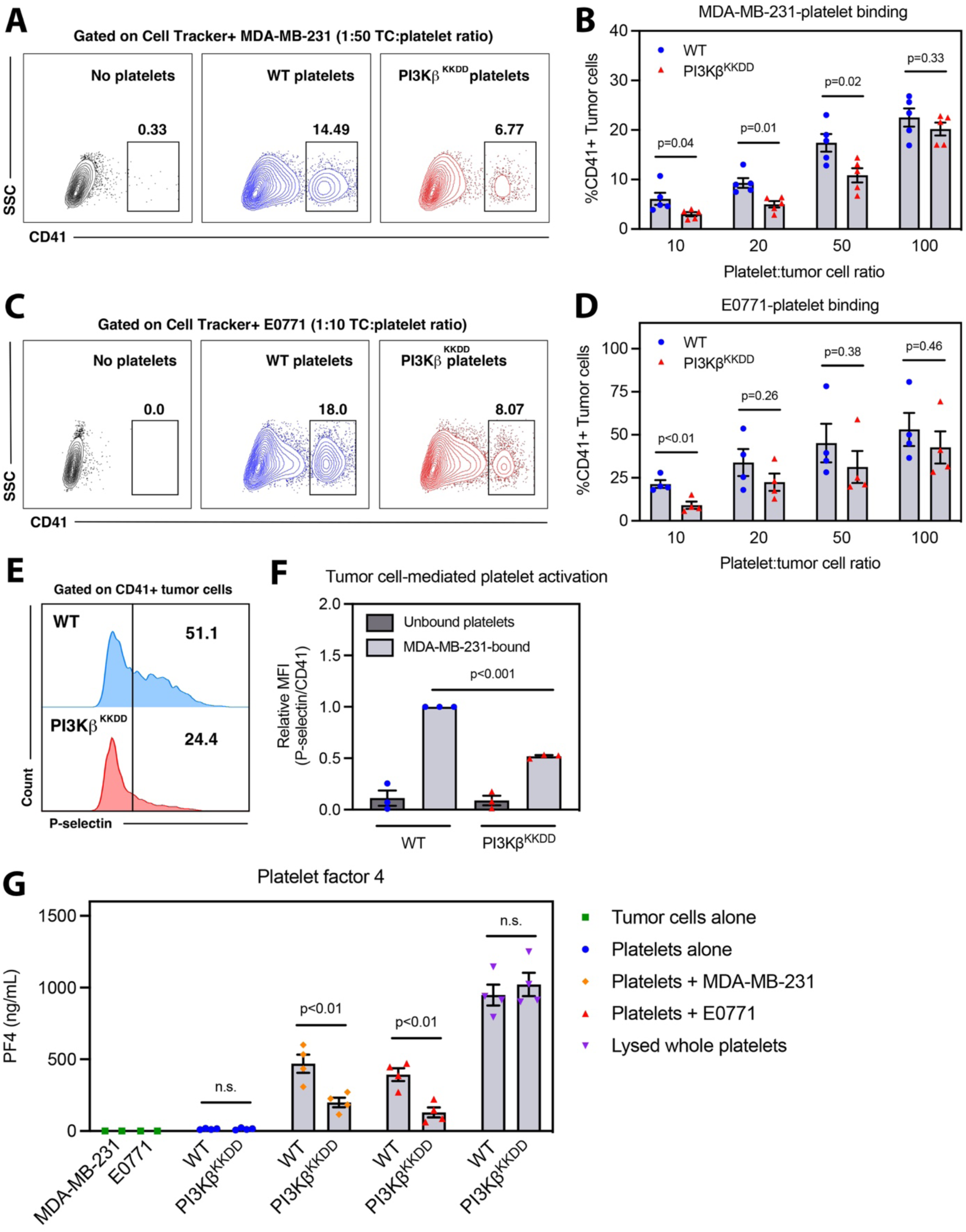
Platelet-tumor cell interactions and platelet activation on tumor cells require platelet PI3Kβ activity. WT or PI3Kβ^KKDD^ platelets and Cell Tracker-labeled MDA-MB-231 or E0771 cells were combined in suspension for 30 min, stained, and analyzed by flow cytometry. FACS plots show platelet interactions with MDA-MB-231 (A) or E0771 (C) cells. (B and D) Quantitation of CD41+ Cell Tracker+ events as a measure of platelet-tumor cell interactions for MDA-MB-231 and E0771 cells, respectively. Data are presented as the mean ± SEM from n=5 (B) or n=4 (D) independent experiments. (E) Analysis and (F) quantitation of relative P-selectin MFI of CD41+ Cell Tracker+ events as a measure of tumor cell-stimulated platelet activation. Data are presented as the mean ± SEM from n=3 independent experiments. (G) ELISA quantitation of Platelet factor 4 release in suspensions of WT or PI3Kβ^KKDD^ platelets with MDA-MB-231 or E0771 tumor cells. Data are presented as the mean ± SEM from n=4 independent experiments.

To confirm these data, we quantified levels of platelet factor 4 (PF4) in the supernatants of platelet-tumor cell suspensions by ELISA, as a measure of platelet degranulation. E0771 and MDA-MB-231 tumor cells stimulated the release of approximately 40-50% of total PF4 in WT platelets (Figure 2G). Activation of PI3Kβ^KKDD^ platelets by tumor cells was significantly impaired, with a greater than two-fold reduction in PF4 release. These data confirm the importance of platelet PI3Kβ activity in mediating platelet activation by tumor cells.

### Platelet PI3Kβ is required for stimulation of Matrigel invasion and EMT-associated morphological changes

Given that we see reduced metastatic potential in tumor cells cultured with PI3Kβ^KKDD^ platelets (Figure 1), we tested the ability of PI3Kβ^KKDD^ platelets to induce an invasive phenotype in tumor cells. We cultured MDA-MB-231 or MCF10A (normal human breast epithelial cells) cells with WT or PI3Kβ^KKDD^ platelets, and then measured tumor cell Matrigel invasion in response to serum. PI3Kβ^KKDD^ platelets were significantly less able to stimulate invasion as compared to WT platelets (Figures 3A and 3B). Similar results were obtained with human platelets pre-treated with the covalent pan-PI3K inhibitor wortmannin (Figure 3C).

**Figure 3.**
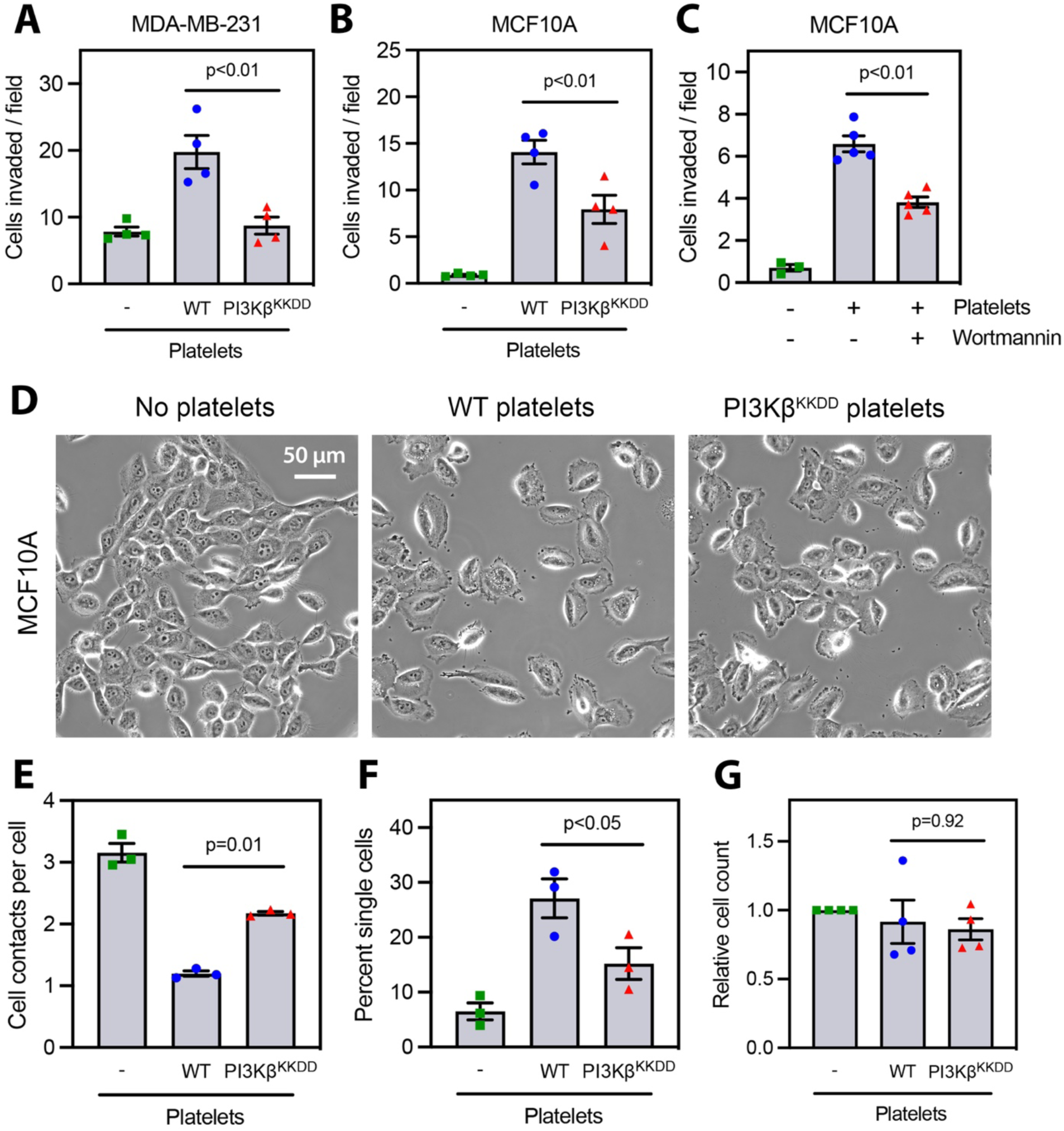
Platelet stimulation of Matrigel invasion and invasion-associated morphological changes requires PI3Kβ. (A-B) MDA-MB-231 or MCF10A cells were cultured for 40 hrs with WT or PI3Kβ^KKDD^platelets and Matrigel invasion was measured after 16 hrs (A) or 20 hrs (B). (C) MCF10A were treated the same as in (B) but were cultured with human platelets pre-treated with Wortmannin or vehicle control. Data are presented as the mean ± SEM from n=4 (A-B) or n=5 (C) independent experiments. (D-G) MCF10A cells were cultured for 40 hrs with WT or PI3Kβ^KKDD^ platelets. (D) Representative 40x phase contrast images from each condition. (E) Number of cell-cell contacts per cell. (F) Percent of cells with no cell-cell contacts. (G) Relative cell counts in each condition. Data are presented as the mean ± SEM from n=3 independent experiments.

Both membrane-bound and soluble factors from platelets stimulate invasion by tumor cells [6, 32, 34]. To test if the defects observed with PI3Kβ^KKDD^ platelets reflect one or both of these fractions, we activated WT and PI3Kβ^KKDD^ platelets with thrombin and then separated the membrane fraction (pellet) from the soluble fraction (releasate). While both the WT platelet pellet and releasate stimulated Matrigel invasion by MCF10A cells, stimulation by the membrane fraction was significantly greater (Figure S3A). However, neither fraction from PI3Kβ^KKDD^ platelets stimulated invasion, suggesting that the defects in PI3Kβ^KKDD^ platelets affect both fractions (Figure S3B).

Platelets induce EMT in tumor cell lines [6]. Co-culture of WT platelets with MCF10A cells results in a reduction in cell-cell contacts, an increase in cell spreading, and the production of extensive lamellipodia (Figures 3D-F). Notably, these responses were attenuated after co-culture with PI3Kβ^KKDD^ platelets. The changes in cell spreading and cell-cell contacts were independent of cell proliferation, which was unaffected by co-culture with WT or mutant platelets (Figure 3G).

### Platelet PI3Kβ is required for stimulation of EMT and PAI-1 expression in breast epithelial and tumor cells

To identify differences in the transcriptional responses of breast epithelial cells co-cultured with WT or PI3Kβ^KKDD^ platelets, we performed RNAseq on MCF10A cells cultured alone or with platelets from three mice of each genotype. We then analyzed the data by Gene Set Enrichment Analysis (GSEA) using the Molecular Signatures Database collection of Hallmark Gene Sets [35, 36]. MCF10A cells cultured with WT platelets were enriched for sixteen gene sets (FDR<0.05) as compared to cells cultured alone (Figure 4A). Consistent with earlier studies [6], the EMT gene set was the most enriched gene set in cells incubated with WT platelets.

**Figure 4.**
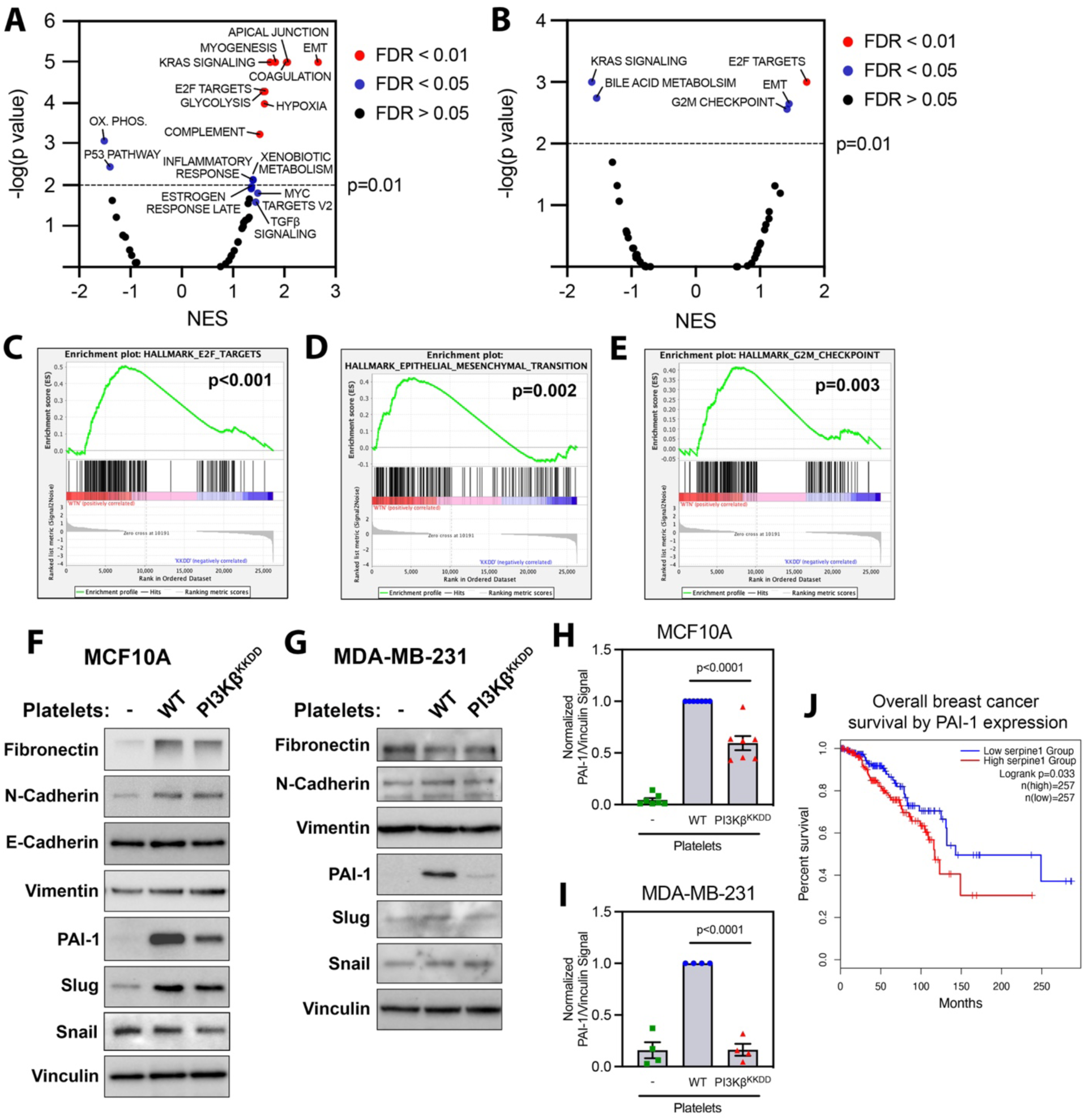
Platelet stimulation of EMT and PAI-1 expression requires PI3Kβ activity. RNAseq data from MCF10A cells cultured for 40 hrs with WT or PI3Kβ^KKDD^ platelets. Gene Set Enrichment Analysis of (A) MCF10A cells cultured with WT platelets vs. MCF10A alone, or (B-E) MCF10A cells cultured with WT vs. PI3Kβ^KKDD^ platelets. (A-B) Analysis of Hallmark Gene Sets, colored by significant enrichment after FDR correction. The mean normalized enrichment score (NES) is displayed on the x-axis. Individual plots demonstrate enrichment of (C) E2F Targets, (D) EMT, and (E) G2M Checkpoint gene sets. MCF10A (F) or MDA-MB-231 (G) cells were cultured for 40 hrs with platelets isolated from WT or PI3Kβ^KKDD^ mice. Lysates were immunoblotted for EMT-associated proteins. (F-G) Representative blots from n=4-7 independent experiments. Expression of PAI-1 in MCF10A (H) or MDA-MB-231 (I) cells was normalized to vinculin. Data are presented as the mean ± SEM from n=7 (H) or n=4 (I) independent experiments. (J) Analysis of overall survival of patients with invasive breast carcinoma from the TCGA database stratified by expression of PAI-1. P-value calculated by logrank test.

When comparing MCF10A cells cultured with WT and PI3Kβ^KKDD^ platelets, the EMT gene set was the second most enriched gene set in cells treated with WT platelets, suggesting an essential role for platelet PI3Kβ in platelet-stimulated EMT (Figure 4B). The other gene sets significantly enriched by co-culture with WT versus PI3Kβ^KKDD^ platelets were E2F targets and G2M checkpoint (Figures 4C-4E). Interestingly, these two gene sets are the most enriched gene sets in metastasis-seeding regions of primary lung tumors [37], consistent with a role for platelet PI3Kβ signaling in metastasis.

To validate the role of platelet PI3Kβ in EMT induction, lysates from MCF10A and MDA-MB-231 co-cultured with WT or PI3Kβ^KKDD^ platelets were immunoblotted for seven EMT-associated proteins previously reported to be stimulated by platelet co-culture [6] (Figures 4F and 4G). While several proteins were induced by platelet co-culture, the only protein that was differentially expressed after co-culture with WT versus PI3Kβ^KKDD^ platelets was PAI-1 (Figures 4H and 4I). PAI-1 is an established regulator of EMT and its expression is associated with poor outcomes in breast cancer patients [38, 39]. Indeed, breast cancer patient data from the TCGA database shows that high expression of PAI-1 is associated with decreased overall survival (Figure 4J). Although platelets express PAI-1, we are confident that our data reflect the induction of PAI-1 in the MCF10A and MDA-MB-231 cells: the antibody used to detect PAI-1 in these experiments is human-specific, and control experiments show that our washing procedure removes platelets from greater than 96-98% of the MCF10A or MDA-MB-231 cells (Figure S4).

### Breast epithelial and tumor cells co-cultured with PI3Kβ^KKDD^ platelets show reduced activation of the TGF-β/Smad and YAP pathways

PAI-1 expression in tumor cells is stimulated by platelet TGF-β released from platelet α-granules [6, 40], as well as by crosstalk between the TGF-β/Smad and YAP pathways [41]. To determine if mutation of PI3Kβ affects α-granules, we isolated WT or PI3Kβ^KKDD^ platelets and analyzed the size and number of granules by transmission electron microscopy (Figure S5A). PI3Kβ^KKDD^ platelets were, on average, 23% larger than WT platelets (Figure S5B) and contained an average of 16% more α-granules than WT platelets (Figure S5C). There was, however, no difference in the size of the granules between genotypes (Figure S5D).

Given these data, it is likely that the reduced induction of tumor cell PAI-1 is due to decreased α-granule secretion and not an inherent α-granule defect. We therefore measured TGF-β release from platelets co-incubated with tumor cells. ELISA data from lysed whole platelets showed no difference in the total or active TGF-β content between WT and PI3Kβ^KKDD^ platelets (Figures 5A and 5B). However, releasate from PI3Kβ^KKDD^ platelet-tumor cell suspensions showed significantly lower levels of both total and active TGF-β as compared to WT platelet suspensions (Figures 5A and 5B). We also examined the signaling pathways upstream of PAI-1 expression, by immunoblotting lysates from co-cultures of MCF10A and MDA-MB-231 with platelets. PI3Kβ^KKDD^ platelets were less able to activate Smad2 (Figures 5B-E) and YAP (Figures 5F-I) in both cell lines. These data suggest that failure of PI3Kβ^KKDD^ platelets to stimulate tumor cell EMT and PAI-1 expression reflects defects in their activation of TGFβ and YAP signaling in target cells.

**Figure 5.**
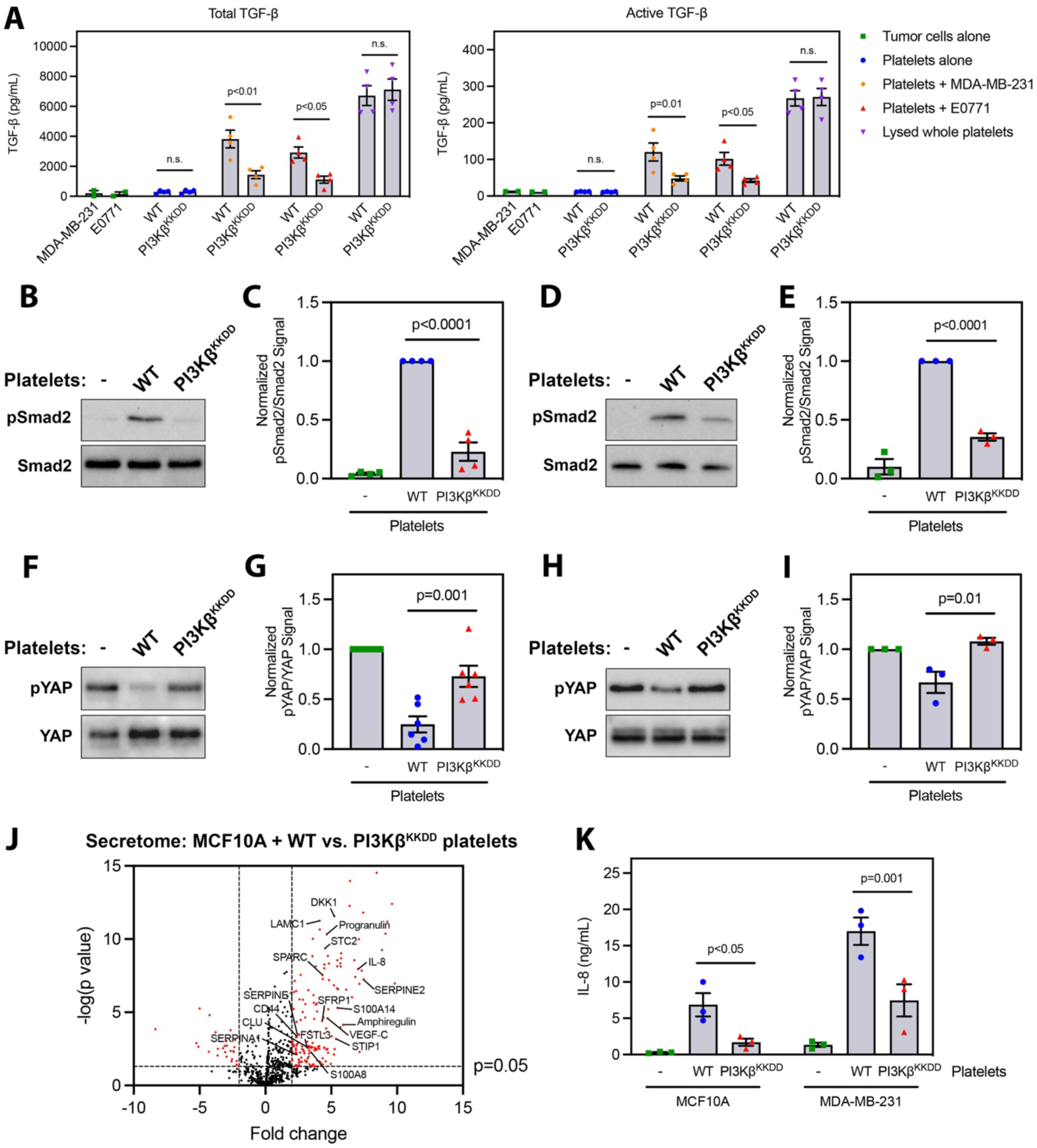
Platelet PI3Kβ activity is required for activation of tumor cell TGF-β/Smad and YAP pathways. (A) ELISA quantitation of murine TGF-β release in suspensions of WT or PI3Kβ^KKDD^ platelets with MDA-MB-231 or E0771 tumor cells. Data are presented as the mean ± SEM from n=4 independent experiments. (B-I) MCF10A (B-C, F-G) or MDA-MB-231 cells (D-E, H-I) cells were cultured for 24 hrs (B-E) or 40 hrs (F-I) with WT or PI3Kβ^KKDD^ platelets. Lysates were immunoblotted for pSmad2/Smad2 (B-E) or pYAP/YAP (F-I). The pSmad2 signal was normalized to total Smad2 in MCF10A (C) or MDA-MB-231 (E) cells. The pYAP signal was normalized to total YAP in MCF10A (G) or MDA-MB-231 (I) cells. Data are presented as the mean ± SEM from n=4 (C), n=3 (E, I), or n=6 (G) independent experiments. (J) Volcano plot of proteins identified in mass spectrometry analysis of conditioned media from MCF10A cells cultured for 12 hrs with WT or PI3Kβ^KKDD^ platelets. Proteins enriched greater than 2-fold with p < 0.05 from WT vs. PI3Kβ^KKDD^ platelet co-cultures are highlighted in red. Proteins known to be involved in cancer or metastasis are labeled. (K) ELISA quantitation of human IL-8 in conditioned medium from cultures of MCF10A or MDA-MB-231 with WT or PI3Kβ^KKDD^ platelets. Data are presented as the mean ± SEM from n=3 independent experiments.

To investigate how downregulation of these pathways affects the secretome of breast epithelial cell cultures, we performed proteomic analysis on conditioned medium from MCF10A cells cultured with WT or PI3Kβ^KKDD^ platelets. Of the 702 proteins identified in the conditioned medium of these co-cultures, significant differences were found in 365 of these proteins. 195 proteins were enriched by at least 2-fold in WT samples (2.0-9.8-fold), while 35 proteins were enriched by at least two-fold in PI3Kβ^KKDD^ samples (2.0-8.4-fold) (Figure 5J). Table 1 highlights eighteen proteins with known roles in cancer or metastasis that showed significant enrichment in the secretomes of MCF10A cultured with WT versus PI3Kβ^KKDD^ platelets. These data identified a 2.4-fold difference in PAI-1 (SERPINE1) levels, as well as a 7.0-fold difference in IL-8 levels. These data were confirmed by ELISA, which showed significant 4.0-fold and 2.3-fold reductions in IL-8 concentrations in MCF10A and MDA-MB-231 cells, respectively, co-cultured with PI3Kβ^KKDD^ platelets (Figure 5K).

**Table 1.** Secretome of MCF10A cells cultured with WT versus PI3Kβ^KKDD^ platelets: cancer and metastasis-associated proteins.

| Name | Function | Fold change | P value |
| --- | --- | --- | --- |
| SERPINE2 | Extracellular serine protease inhibitor; breast cancer metastasis [54] | 7.5 | <0.0001 |
| IL-8 | Chemokine; Platelet-mediated metastasis; immune cell recruitment [32, 53] | 7.0 | <0.0001 |
| Amphiregulin | EGFR ligand; osteosarcoma and gastric cancer metastasis [50, 51] | 5.9 | <0.0001 |
| S100A14 | Calcium binding protein; Breast cancer invasion and metastasis [83, 84] | 5.5 | <0.0001 |
| STIP1 | HSP co-chaperone; Secreted STIP1 promotes cancer proliferation [85] | 5.4 | 0.001 |
| DKK1 | Secreted Wnt signaling inhibitor; Breast cancer metastasis [86] | 5.2 | <0.0001 |
| VEGF-C | Growth factor; Breast cancer cell proliferation, invasion, and metastasis [52] | 4.7 | <0.0001 |
| Progranulin | Secreted glycoprotein; tumor immune evasion [87] | 4.7 | <0.0001 |
| STC2 | Secreted glycoprotein; HNSCC metastasis; Glioblastoma malignant transformation [88, 89] | 4.5 | <0.0001 |
| SPARC | Matrix-associated glycoprotein; Renal cell carcinoma metastasis [90] | 4.3 | <0.0001 |
| SFRP1 | Secreted modulator of Wnt signaling; Renal cell carcinoma invasion and metastasis [91] | 4.3 | <0.0001 |
| LAMC1 | Laminin subunit; Hepatocellular carcinoma prognosis, cell proliferation, and invasion [92] | 4.1 | <0.0001 |
| S100A8 | Calcium binding protein; Colorectal cancer invasion, EMT, and metastasis [61] | 3.5 | 0.006 |
| FSTL3 | Secreted glycoprotein; Colorectal cancer invasion, EMT, and metastasis [69] | 3.2 | 0.003 |
| Clusterin | Extracellular molecular chaperone; Breast and prostate cancer EMT and metastasis [62, 93] | 3.1 | 0.002 |
| SERPINE1 | Extracellular serine protease inhibitor; Breast cancer invasion, EMT, and metastasis [38-40] | 2.4 | 0.0003 |
| CD44 | Cell surface and secreted glycoprotein; Breast cancer stem cell marker [94] | 2.4 | 0.0004 |
| SERPINA1 | Extracellular serine protease inhibitor; Colorectal cancer invasion, EMT, and metastasis [55] | 2.4 | 0.007 |

### Activation of platelets by tumor cells is reduced in PI3Kβ^KKDD^ mice

PI3Kβ^KKDD^ platelets show reduced activation by tumor cells *in vitro* (Figures 2E-G). To test if this also occurs *in vivo*, blood from WT or PI3Kβ^KKDD^ mice bearing 1.5 cm^3^ orthotopic GFP-E0771 tumors was analyzed by flow cytometry (Figure S6). PI3Kβ^KKDD^ mice showed a modest trend toward reduced CTCs (Figure 6A), but we saw no significant differences in *in vivo* platelet-CTC binding in WT versus PI3Kβ^KKDD^ mice (Figure 6B). This likely reflects that fact that platelet: tumor cell ratios are much higher *in vivo* than in our *in vitro* experiments (Figures 2B and 2D). However, the activation of platelets bound to CTCs *in vivo* was significantly reduced in PI3Kβ^KKDD^ mice (Figure 6C), which directly correlates with our *in vitro* data showing defective activation of PI3Kβ^KKDD^ platelets by tumor cells (Figure 2E-G).

**Figure 6.**
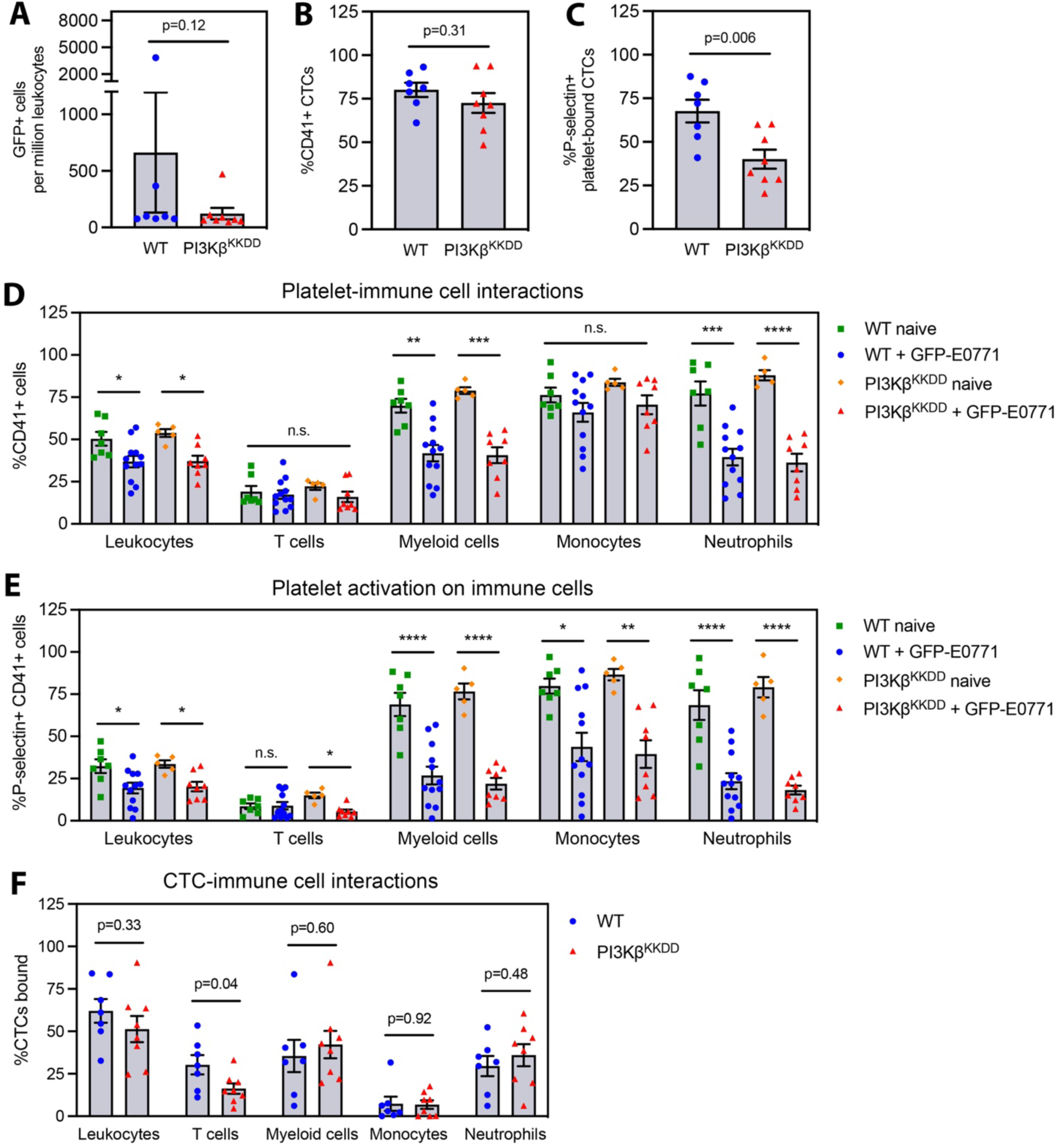
*In vivo* activation of platelets by tumor cells is reduced in PI3Kβ^KKDD^ mice. Blood from WT or PI3Kβ^KKDD^ mice, either naïve or bearing orthotopic GFP-E0771 tumors, was collected by cardiac puncture and analyzed by flow cytometry. (A) GFP+ circulating tumor cells (CTC) normalized to CD45+ leukocytes. (B) Platelet-CTC interactions expressed as the percent of GFP+ CTCs that stained positive for CD41. (C) Tumor cell-stimulated platelet activation, defined as the percent of GFP+CD41+ CTCs that stained positive for P-selectin. (D) Platelet-immune cell interactions, expressed as the percent of each immune cell type that stained positive for CD41. (E) Immune cell-stimulated platelet activation, defined as the percent of CD41+ immune cells that stained positive for P-selectin. (F) CTC-immune cell interactions, expressed as the percent of GFP+ CTCs that stained positive for markers of individual immune cell types. Data are presented as the mean ± SEM from 7-8 (A-C, F) or 5-12 (D-E) mice per group. *p<0.05; **p<0.01, ***p<0.001, ****p<0.0001 by ordinary one-way ANOVA.

Platelet PI3Kβ is required for platelet-leukocyte interactions and platelet-mediated activation of innate immune cells in the context of infection [18]. Given the complex modulation of the immune system in cancer, we analyzed platelet-leukocyte interactions and activation of platelets bound to immune cells in naïve and tumor-bearing WT and PI3Kβ^KKDD^ mice. Interestingly, the presence of an orthotopic GFP-E0771 mammary tumor caused a decrease in platelet-immune cell interactions and activation of platelets bound to immune cells in mice of both genotypes (Figures 6D-E). These differences were not due to changes in platelet or immune cell numbers in the tumor-bearing mice (Figures S7A-F), although tumor-bearing mice did exhibit a significant anemia in both genotypes (Figures S7G-I). Furthermore, we saw no significant differences in either platelet binding or activation on immune cells in PI3Kβ^KKDD^ mice as compared to control mice, in either naïve or tumor-bearing mice (Figures 6D-E). Taken together, these data suggest that mutation of platelet PI3Kβ causes a selective disruption of platelet activation by tumor cells but does not affect their activation by immune cells.

Since it is known that platelets mediate recruitment of neutrophils to CTCs [1], we tested whether stromal PI3Kβ activity is required for CTC-immune cell interactions in tumor-bearing mice. We did not see any effect of stromal PI3Kβ activity on CTC-neutrophil interactions (Figure 6F). However, we did find a significant reduction in T cell binding to CTCs in PI3Kβ^KKDD^ mice (Figure 6F). The mouse used for these studies is a whole-body knock-in, and this effect could be due to altered PI3Kβ signaling in T cells, platelets, or another intermediary cell type. Future studies will explore the cell types involved, and test whether decreased CTC binding to T cells has an effect on metastasis.

Our data describe a role for platelet PI3Kβ in regulating tumor cell invasion, epithelial-mesenchymal transition, and metastasis (Figure 7). Mutation of platelet PI3Kβ abrogates these phenotypes, suggesting a broadened clinical application of PI3Kβ-selective inhibitors to treat metastasis.

**Figure 7.**
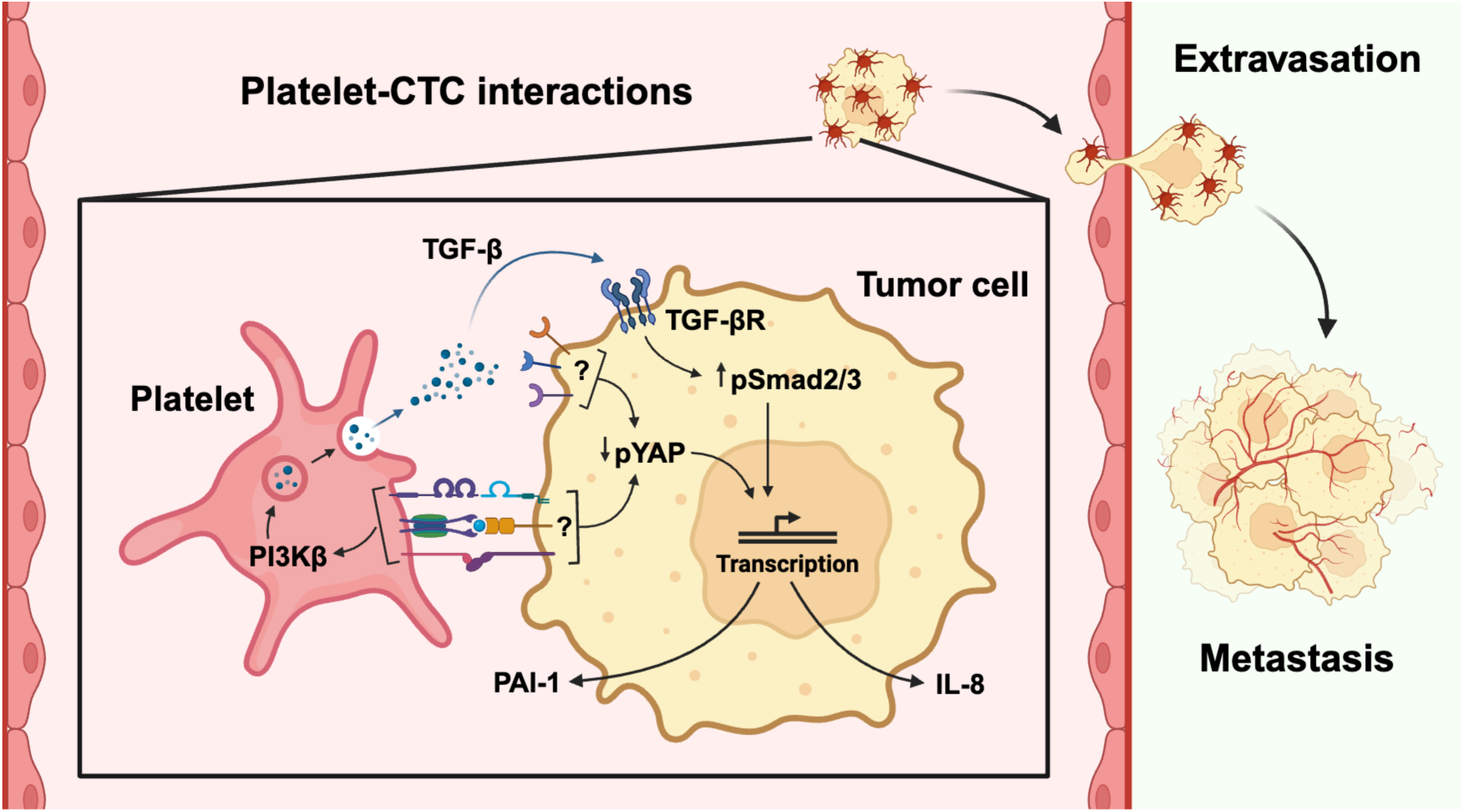
Platelet PI3Kβ regulates platelet-mediated tumor cell invasion, EMT, and metastasis. Platelets bind to circulating tumor cells in the bloodstream. This interaction results in activation of platelets, dependent on PI3Kβ. Platelet activation results in release of α-granule contents, including TGF-β. PI3Kβ-dependent release of TGF-β and other ligands acts on the tumor cell to stimulate Smad and YAP signaling. These signaling pathways regulate transcription of genes associated with EMT and invasion, including PAI-1 and IL-8. As a result of this altered transcriptomic state, tumor cells have increased metastatic potential. Mutation of platelet PI3Kβ abrogates these phenotypes.

## Discussion

We have previously shown that PI3Kβ is a critical regulator of tumor cell invasion [22–24]. In the current study, we focus on the role of stromal PI3Kβ in tumor metastasis. We produced a novel knock-in mouse, which expresses a KK^526/527^DD substitution in the Gβγ binding site of *PIK3CB* (PI3Kβ^KKDD^). This mutation blocks PI3Kβ activation but does not diminish its basal activity [20], and PI3Kβ^KKDD^ knock-in mice are viable and fertile. This contrasts with *PIK3CB* kinase-dead knock-in mice, which exhibit increased embryonic lethality [42]. Our new mouse model has allowed us to examine the role of stromal PI3Kβ in tumor growth and metastasis.

The growth of orthotopic syngeneic breast tumors in PI3Kβ^KKDD^ mice was identical to that seen in control mice. However, we observed a strong trend toward a reduction in spontaneous metastasis to the lung. These data support the hypothesis that PI3Kβ plays an important role in breast cancer metastasis in both tumor cells [22–24] and stromal cells (this study). The global PI3Kβ^KKDD^ knock-in affects multiple cell types involved in metastasis. In this study, we focused on the role of pro-metastatic signaling by platelets, as these cells are known to depend heavily on PI3Kβ signaling for many responses [10–14].

Incubation of tumor cells with wild type platelets increased experimental metastasis and led to morphological changes consistent with EMT, as well as increases in the TGF-β target gene PAI-1 and the stemness marker Slug. Tumor cell PAI-1 expression is regulated by crosstalk between the TGF-β/Smad and YAP pathways [41], both of which were activated by co-culture with platelets. However, platelet-treated cells still expressed the epithelial marker E-cadherin, suggesting the induction of a hybrid E/M state [43], known to be highly tumorigenic and metastatic [44, 45].

Importantly, these responses were all attenuated in tumor cells incubated with PI3Kβ^KKDD^ platelets. Consistent with these biochemical analyses, GSEA of RNAseq data from breast epithelial cells co-cultured with wild type versus PI3Kβ^KKDD^ platelets showed significant differences in just three gene sets: EMT, E2F targets, and G2M checkpoint. While we did not investigate the activity of G2M and E2F pathways in tumor cells co-cultured with PI3Kβ^KKDD^ platelets, these pathways are markers of breast cancer invasiveness and are predictive of chemotherapeutic response [46, 47]. Interestingly, they are also the two most highly enriched gene sets in metastasis-seeding regions of primary lung tumors [37]. There have also been reports of crosstalk between E2F and TGF-β/Smad pathways [48, 49], which may relate to the downregulation of both pathways that we observe in this study.

Proteomic analysis of secretomes from MCF10A-platelet co-cultures identified several pro-metastatic factors whose secretion depends on platelet PI3Kβ. These include ligands such as amphiregulin [50, 51], VEGF-C [52], and IL-8 [32]. IL-8 secretion by tumor cells is known to be stimulated by platelet co-culture and is associated with breast cancer invasion and metastasis [32]. IL-8 also plays an important role in the recruitment of neutrophils and myeloid-derived suppressor cells to primary tumors [53]. The decreased IL-8 induction in MCF10A-PI3Kβ^KKDD^ platelet co-cultures was confirmed by ELISA. Co-culture with WT but not PI3Kβ^KKDD^ platelets also led to increases in several Serpin family protease inhibitors, including SERPINE1 (PAI-1, discussed above), SERPINE2, and SERPINA1, all known to promote metastasis [38–40, 54, 55]. Numerous proteins enriched in WT platelet co-cultures are regulated by TGF-β signaling, including SERPINE2 [56], IL-8 [57], Amphiregulin [58], VEGF-C [59], LAMC1 [60], S100A8 [61],

Clusterin [62], SERPINE1 [41], and CD44 [63]. Similarly, we detected several proteins regulated downstream of YAP, including IL-8 [64], Amphiregulin [65], DKK1 [66], VEGF-C [67], S100A8 [68], FSTL3 [69], and SERPINE1 [41], as well as the stemness marker CD44 and the matricellular protein SPARC, which activate YAP signaling [70, 71]. Together, these data show that platelet PI3Kβ is required for the stimulation of a pro-metastatic secretory response in tumor cells, due in part to stimulation of TGF-β and YAP signaling.

The mechanism of decreased pro-metastatic signaling by PI3Kβ^KKDD^ platelets is likely due to their decreased activation when bound to tumor cells, as P-selectin surface exposure on tumor cell-bound platelets was decreased both *in vitro* and *in vivo*. P-selectin exposure reflects secretion of α-granules, which contain TGF-β, PF4, PDGF and other ligands. Consistent with decreased α-granule secretion, tumor cell-stimulated release of PF4 and platelet TGF-β was 2 to 3-fold lower in PI3Kβ^KKDD^ platelets. While we do not know how platelets stimulate the YAP pathway in tumor cells, α-granule contents such as thromboxane A2 and PDGF activate YAP in other systems [72, 73]. Several studies have shown that knockout, knockdown, or pharmacologic inhibition of YAP reduces TGF-β-stimulated Smad2/3 nuclear translocation and transcription of target genes [74–76]. The combination of reduced local TGF-β release and impaired activation of YAP signaling likely contributes to the loss of pro-metastatic signaling and hybrid E/M induction that we see in PI3Kβ^KKDD^ platelet-tumor cell co-cultures.

An unexpected finding from this study was a PI3Kβ-independent decrease in platelet-immune cell interactions in tumor-bearing mice. The activation of platelets by immune cells results in activation of innate immune cells and upregulation of the immune response [77]. A reduction in this response may reflect an immunosuppressive environment produced by factors secreted from the primary tumor or CTCs. However, in contrast to the reported role of platelet PI3Kβ in regulating platelet-leukocyte interactions and platelet-mediated activation of innate immune cells in the context of bacterial pneumonia [18], we found no role for platelet PI3Kβ in regulating these interactions in the context of cancer.

While platelet-immune cell interactions did not depend on stromal PI3Kβ activity, T cell binding to CTCs was decreased in PI3Kβ^KKDD^ mice. It is possible that mutation of platelet PI3Kβ impacts T cell-CTC interactions, however, it may be more likely that PI3Kβ in T cells is involved. The predominant PI3Ks in T cells are PI3Kγ and PI3Kδ; PI3Kγ acts largely downstream of T cell GPCRs while PI3Kδ is essential for TCR signaling, integrin activation, and mediating interactions with other cells [78]. The role of T cell PI3Kβ is not well understood. Our data may suggest a potential unappreciated role for PI3Kβ signaling in T cell signaling.

Isoform selective PI3Kβ inhibitors were originally developed as anti-thrombotic agents [10, 79, 80]. More recently, these inhibitors have been explored as anti-cancer agents [81]. However, clinical trials have focused on treatment of PTEN-deficient cancers, which are known to depend on PI3Kβ activity [81, 82]. Our study demonstrates that stromal PI3Kβ regulates breast cancer metastasis and reveals a novel mechanism by which platelet PI3Kβ contributes to this process. This work, along with our work on PI3Kβ signaling in tumor cells [22–24], suggests that isoform selective PI3Kβ inhibitors may have unappreciated utility in the treatment of metastasis in solid tumors.

## Acknowledgements

The authors thank the following contributors and shared resources at the Montefiore Einstein Comprehensive Cancer Center (MECCC) for their help: the Genomics Shared Resource for analysis of RNAseq data, the Analytical Imaging Facility for transmission electron microscopy data, the Animal Model Shared Resource for maintenance of mouse strains and for assistance with tail vein injections, the Structural Biology Shared Resource for acquisition and analysis of mass spectrometry data, and the Flow Cytometry Shared Resource for acquisition and analysis of flow cytometry data.

This work was supported by National Institutes of Health (NIH) National Cancer Institute grants: F30CA265236, P01CA257885, and P30CA013330. It was also supported by NIH National Institute of General Medical Sciences grant T32GM149364 (Medical Scientist Training Program) and NIH Office of the Director grant 1S10OD026833 (shared instrument grant). The Sidoli lab gratefully acknowledges for funding the Hevolution Foundation (AFAR), Einstein-Mount Sinai Diabetes center, Merck, Relay Therapeutics, Deerfield (Xseed award) and the NIH Office of the Director (S10OD030286).

## Author Contributions

R.C.G. designed the research, acquired and analyzed the data, and wrote the manuscript; A.H. and M.J.G. helped design, acquire, and analyze RNAseq research and data; J.T.A. and S.Sidoli helped acquire and analyze mass spectrometry data; A.L. helped acquire and analyze *in vivo* data; J.Z. and S.A.P. helped design flow cytometry-based research and analyze data; W.Y., M.R.G, and S. Seshagiri designed and produced the PI3Kβ^KKDD^ knock-in mouse model; and A.R.B. and J.M.B designed and supervised the research, contributed to discussion, and edited the manuscript.

## Data Availability Statement

The data generated in this study are available upon request from the corresponding authors. The mass spectrometry proteomics data have been deposited to the ProteomeXchange Consortium via the PRIDE partner repository with the dataset identifier PXD050856.

For reviewers: **Username:**; **Password:** 6lPI1pam

