## Supplemental methods and figures for "Platelet PI3Kβ regulates breast cancer metastasis"

**Cell culture**

MDA-MB-231 cells (ATCC) were maintained in DMEM (Gibco; 11965092) with 10% fetal bovine serum (FBS) (R&D Systems; S11550) and 1 mM sodium pyruvate (Cytiva; SH30239.01). MCF10A cells (ATCC) were maintained in DMEM/F12 (Cytiva; SH30023FS) supplemented with 5% horse serum (Gibco; 26050088), 20 ng/mL EGF (Millipore Sigma; 01107MI), 0.5 μg/mL hydrocortisone (Thermo Scientific Chemicals; AC352450010), 10μg/mL insulin (Gibco; 12585014), and 100 ng/mL cholera toxin (Millipore Sigma; C8052). E0771 cells were obtained from CH3 BioSystems, and GFP-E0711 cells were provided by Dr. John Condeelis (Albert Einstein College of Medicine). Both lines were maintained in RPMI 1640 with 10% FBS, 1 mM sodium pyruvate, and 20 mM HEPES (Fisher Scientific; ICN1688449). The Cerulean-PyMT cell line was provided by Dr. David Entenberg (Albert Einstein College of Medicine) and maintained in DMEM with 10% FBS and 1 mM sodium pyruvate. All cell lines were confirmed to be free of mycoplasma (ATCC; 30-1012K).

**Enzyme-linked immunosorbent assays**

To analyze PF4 and TGF-β release, platelets alone, tumor cells alone, or tumor cells and platelets together were incubated in PBS for 30 min at 25^o^C with rotation as above. Supernatants were collected by centrifugation. Alternatively, platelet lysates were collected after sonicating platelets in PBS. PF4 and TGF-β concentrations were determined using the mouse CXCL4/PF4 or mouse TGF-β1 DuoSet ELISA Kits (R&D Systems; DY595 and DY1679-05, respectively) according to the manufacturer’s instructions. For IL-8 analysis, conditioned medium was collected from 40-hour platelet-tumor cell co-cultures and sterile filtered with a 0.20 µm pore syringe filter (Corning; 431229). IL-8 concentrations were determined using the human IL-8/CXCL8 DuoSet ELISA Kit (R&D Systems; DY208) according to the manufacturer’s instructions.

**Matrigel invasion**

Tumor cells were co-cultured for 40 hours without or with platelets isolated from WT or PI3Kβ^KKDD^ mice as described above. For experiments with platelet pellets and releasate, platelets were stimulated with 0.1 U/mL thrombin (Millipore Sigma; 605195) in MTHB for 10 min at 37^o^C and pelleted at 8,000xg for 8 min. The supernatant was used as the “platelet releasate” and the pellet was resuspended in an equal volume of MTHB and used as the “platelet pellet.” MCF10A were incubated with platelets, pellets, or releasate for 40 hours, washed in PBS and detached with trypsin. Cells were washed in PBS and resuspended in DMEM with 1% FBS. Growth factor-reduced Matrigel-coated transwell inserts (Corning; 354483) were inserted into wells containing DMEM with 10% FBS (lower chamber). 2.5x10^4^ tumor cells were transferred to the upper chamber of the transwell inserts and allowed to invade for 16 or 20 hours (MDA-MB-231 and MCF10A, respectively). Cells remaining on the upper surface of the insert were removed using a cotton tipped applicator, and cells on the lower surface of the insert were fixed with 4% formaldehyde for 15 min. The inserts were removed and mounted onto slides with DAPI Fluoromount-G (Southern Biotech; 0100-20). The entire underside of the insert was imaged using a Nikon Eclipse E400 upright microscope equipped with a Nikon 10x (0.25 N.A.) objective. ImageJ was used for automated counting of nuclei by thresholding.

**Cell morphology**

MCF10A cells were seeded onto MatTek dishes (Fisher Scientific; NC9268399) and cultured without or with platelets for 40 hours, and then washed three times to remove platelets. Live cells were imaged using an Olympus IX70 inverted microscope equipped with a UPlanFl 40x (0.75 N.A.) objective. The number of cell-cell contacts per cell were counted (at least 100 cells per condition). Alternatively, the number of isolated cells (no cell-cell contacts) were counted and expressed as a percentage of the total cells in the field.

**Western blotting**

MCF10A or MDA-MB-231 cells were cultured for 40 hours without or with platelets isolated from WT or PI3Kβ^KKDD^ mice as described above. Cells were washed three times to remove platelets and lysed in Laemmli buffer lacking bromophenol blue or BME and supplemented with 1 mM PMSF, 5 μg/mL leupeptin, 10 μg/mL pepstatin A, 5 μg/mL aprotinin (Millipore Sigma; L9783, 516481, and A6106, respectively), 1 mM DTT, and Phosphatase Inhibitor Cocktails 2 and 3 (Sigma; P5726 and P0044). Cell lysates were sonicated and then boiled for 5 min. Protein concentrations were determined by DC protein assay (BioRad; 5000116). Lysates were mixed with sample buffer and DTT to a final concentration of 100 mM. Samples (30-100 μg protein) were separated by SDS-PAGE and blotted using antibodies against: N-Cadherin (BD Biosciences; 610920), Vinculin (Santa Cruz; sc-73614), Fibronectin (Abcam; ab2413), E-Cadherin (Cell Signaling Technologies (CST); 3195), Snail (CST; 3879), Slug (CST; 9585), PAI-1 (CST; 49536), Smad2 (CST; 5339), pSer^465/467^Smad2 (CST; 3108), YAP (CST; 14074), pSer^127^YAP (CST; 13008), and Vimentin (CST; 5741). Blots were developed using the Super Signal West Pico PLUS Chemiluminescent Substrate (Thermo Scientific; 34580) and imaged with a Biorad Chemidoc imaging station. Bands were quantified by densitometry using ImageJ software.

**Platelet spreading**

Glass coverslips were coated with 0.01% poly-L-lysine (Millipore Sigma; P4707) for 15 min at 25^o^C, washed with PBS, and coated with 200 μg/mL fibrinogen (Millipore Sigma; F3879) in PBS for 1 hour at 37^o^C. Platelets isolated from WT or PI3Kβ^KKDD^ mice were resuspended in MTHB supplemented with 2 mM calcium. 5x10^6^ platelets were added to each coverslip and allowed to attach and spread for 30 min at 37^o^C. Coverslips were fixed in 4% formaldehyde for 15 min, permeabilized with 0.1% NP-40 for 10 min, and stained with Rhodamine phalloidin for 30 min (Invitrogen; R415). Mounted coverslips were imaged using a Nikon Eclipse E400 upright microscope equipped with a Nikon 100x (1.25 N.A.) objective. ImageJ software was used to determine platelet area by thresholding images to highlight Rhodamine positive platelets.

**LC-MS/MS Sample preparation**

MCF10A cells were cultured in serum-free medium for 12 hours without or with WT or PI3Kβ^KKDD^ platelets as described above. Conditioned medium was collected and centrifuged at 30,000xg for 15 min at 4^o^C to remove cells and cell debris. Samples were vacuum dried overnight, solubilized in 5% SDS in 50mM TEAB, reduced with 20 mM DTT at 56^o^C for 30 min, and alkylated with 40 mM iodoacetamide for 30 min at 25^o^C. Phosphoric acid was added to a final concentration of 1.2%. S-trap binding buffer (90% methanol, 10 mM ammonium bicarbonate) was added before loading samples onto S-trap columns (Protifi). Columns were washed with S-trap binding buffer three times and samples were digested with 2 μg trypsin at 47^o^C for one hour. Peptides were eluted and final samples were vacuum dried overnight.

Prior to mass spectrometry analysis, samples were desalted using a 96-well plate filter (Orochem) packed with 1 mg of Oasis HLB C-18 resin (Waters). Briefly, the samples were resuspended in 100 µl of 0.1% TFA and loaded onto the HLB resin, which was previously equilibrated using 100 µl of the same buffer. After washing with 100 µl of 0.1% TFA, the samples were eluted with a buffer containing 70 µl of 60% acetonitrile and 0.1% TFA and then dried in a vacuum centrifuge.

**LC-MS/MS Acquisition and Analysis**

Samples were resuspended in 10 µl of 0.1% TFA and loaded onto a Dionex RSLC Ultimate 300 coupled online with an Orbitrap Fusion Lumos (Thermo Scientific). Chromatographic separation was performed with a two-column system, consisting of a C-18 trap cartridge (300 µm ID, 5 mm length) and a picofrit analytical column (75 µm ID, 25 cm length) packed in-house with reversed-phase Repro-Sil Pur C18-AQ 3 µm resin. Peptides were separated using a 60 min gradient from 4-30% buffer B (buffer A: 0.1% formic acid, buffer B: 80% acetonitrile + 0.1% formic acid) at a flow rate of 300 nL/min. The mass spectrometer was set to acquire spectra in a data-dependent acquisition (DDA) mode. Briefly, the full MS scan was set to 300-1200 m/z in the orbitrap with a resolution of 120,000 (at 200 m/z) and an AGC target of 5x10e5. MS/MS was performed in the ion trap using the top speed mode (2 secs), an AGC target of 1x10e4 and an HCD collision energy of 35. Proteome raw files were searched using Proteome Discoverer software (v2.5, Thermo Scientific) using SEQUEST search engine and the SwissProt human database. The search for total proteome included variable modification of N-terminal acetylation, and fixed modification of carbamidomethyl cysteine. Trypsin was specified as the digestive enzyme with up to 2 missed cleavages allowed. Mass tolerance was set to 10 pm for precursor ions and 0.2 Da for product ions. Peptide and protein false discovery rate was set to 1%. Following the search, data was processed as previously described [1]. Briefly, proteins were log2 transformed, normalized by the average value of each sample and missing values were imputed using a normal distribution 2 standard deviations lower than the mean. Statistical regulation was assessed using heteroscedastic T-test (if p-value < 0.05). Data distribution was assumed to be normal, but this was not formally tested. The mass spectrometry proteomics data have been deposited to the ProteomeXchange Consortium via the PRIDE [2] partner repository with the dataset identifier PXD050856.

**Transmission electron microscopy**

WT or PI3Kβ^KKDD^ platelets were pelleted and fixed with 2.5% glutaraldehyde in 0.1 M sodium cacodylate buffer, postfixed with 1% osmium tetroxide followed by 2% uranyl acetate, dehydrated through a graded series of ethanol, and embedded in LX112 resin (LADD Research Industries, Burlington VT).  Ultrathin (80 nm) sections were cut on a Leica EM Ultracut UC7, stained with uranyl acetate followed by lead citrate, and imaged on a JEOL 1400 Plus transmission electron microscope at 120kv.

**Tumor cell-stimulated thrombocytopenia**

Blood was collected from the submandibular veins of 10–16-week-old WT or PI3Kβ^KKDD^ mice directly into K_2_EDTA-coated blood collection tubes (Greiner Bio-One; 450480). Platelet counts were determined with a hematology analyzer (Genesis, Oxford Sciences). E0771 cells were then detached with trypsin and washed three times with PBS. A total of 5.0x10^5^ cells in 200 μL PBS were injected into the tail veins of the same mice. Three hours later, blood was collected again and platelet counts were determined by hematology analyzer.

**Supplemental Figures and Legends**

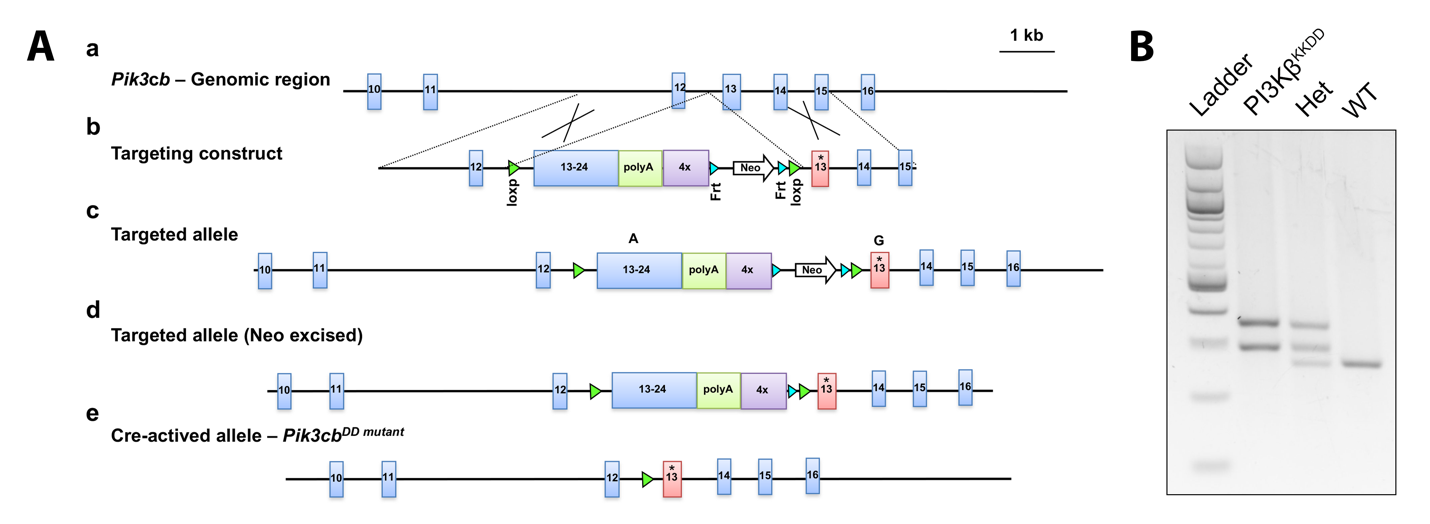

**Supplemental Figure 1. Generation of PI3Kβ^KKDD^ mice.** (A) a. Map of the *PIK3CB* genomic locus with coding exons 10-16 shown in blue. b. Targeting vector used to modify the *PIK3CB* locus. Homology arms for targeting of C57Bl6/N ES cells are indicated. An asterisk indicates the KKDD mutation in exon 13 (pink box). c. Targeted *PIK3CB* locus with the neomycin (Neo) selection marker present. d. PIK3CB^KKDD^ conditional knock-in after FLP-mediate excision of *Neo*. e.  *PIK3CB^KKDD^* knock-in allele after Cre-mediated excision of the floxed wild type *PIK3CB* cassette. (B) PCR genotyping of the knock-in allele.

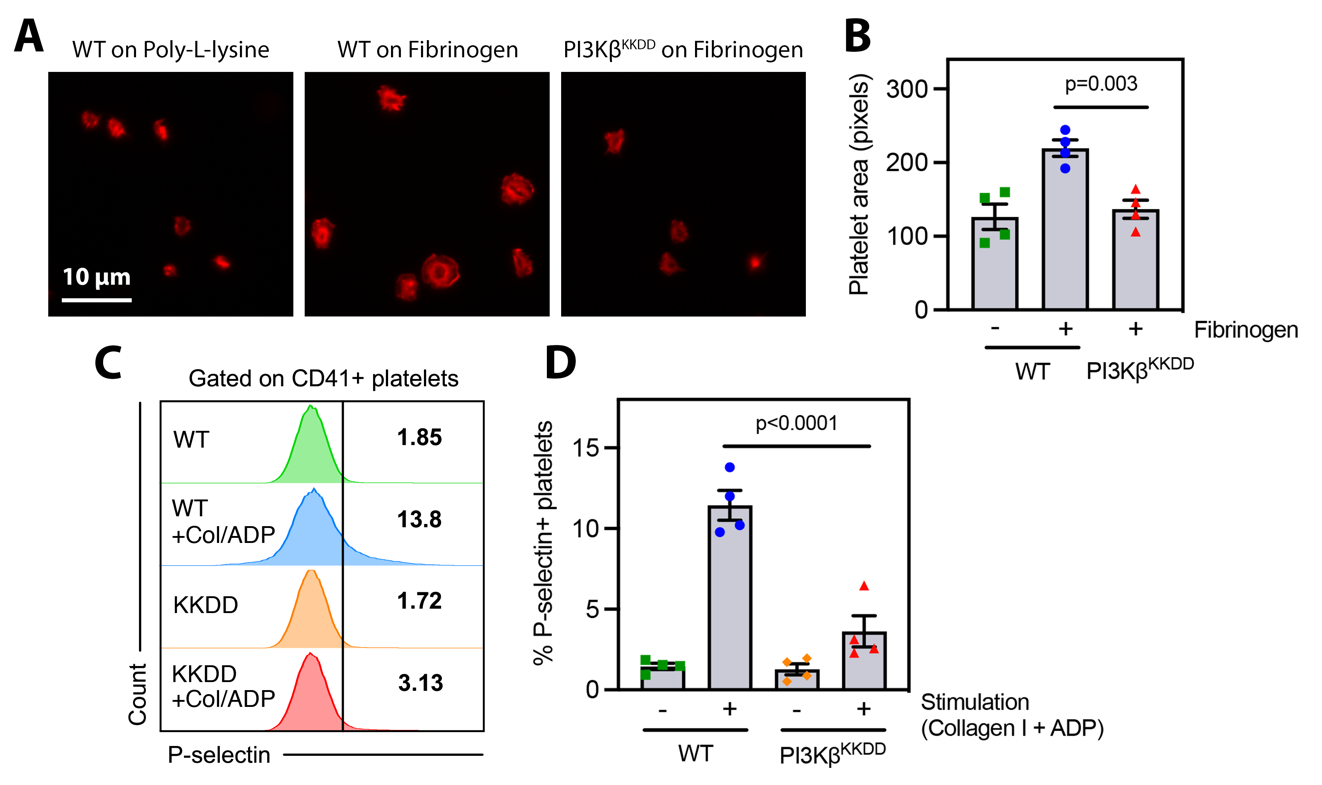

**Supplemental Figure 2.** **PI3Kβ^KKDD^ platelets exhibit decreased spreading and activation.** (A) WT or PI3Kβ^KKDD^ platelets were incubated on poly-L-lysine or fibrinogen coated coverslips for 30 min before imaging. Representative fluorescent images of rhodamine-phalloidin stained platelets are shown for each condition. (B) Quantitation of platelet area. Data are presented as the mean ± SEM from n=4 independent experiments. (C) WT or PI3Kβ^KKDD^ platelets were incubated in suspension with soluble collagen I and ADP for 30 min, stained, and analyzed by flow cytometry. FACS plots show platelet activation for each condition, measured by percent of platelets that stain positive for P-selectin. (D) Quantitation of the percent of P-selectin+ platelets. Data are presented as the mean ± SEM from n=4 independent experiments.

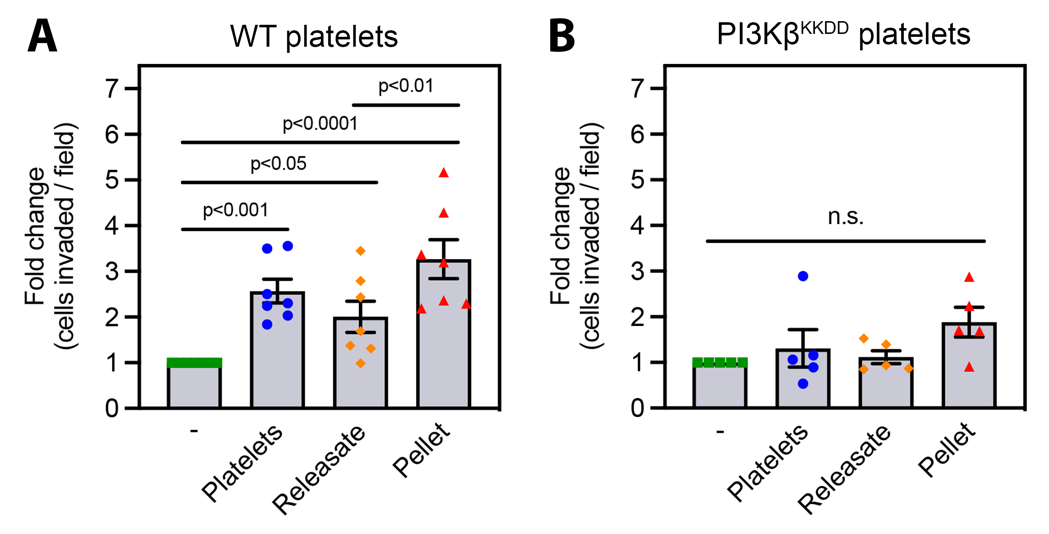

**Supplemental Figure 3. Mutation of platelet PI3Kβ impairs both platelet releasate and platelet pellet stimulation of tumor cell Matrigel invasion.** MCF10A cells were cultured for 40 hrs with either whole platelets, the membrane fraction (pellet) from activated platelets or the soluble fraction (releasate) from activated platelets isolated from WT (A) or PI3Kβ^KKDD^ (B) mice. Matrigel invasion was measured after 20 hrs. Data are presented as the mean ± SEM from n=7 (A) or n=5 (B) independent experiments.

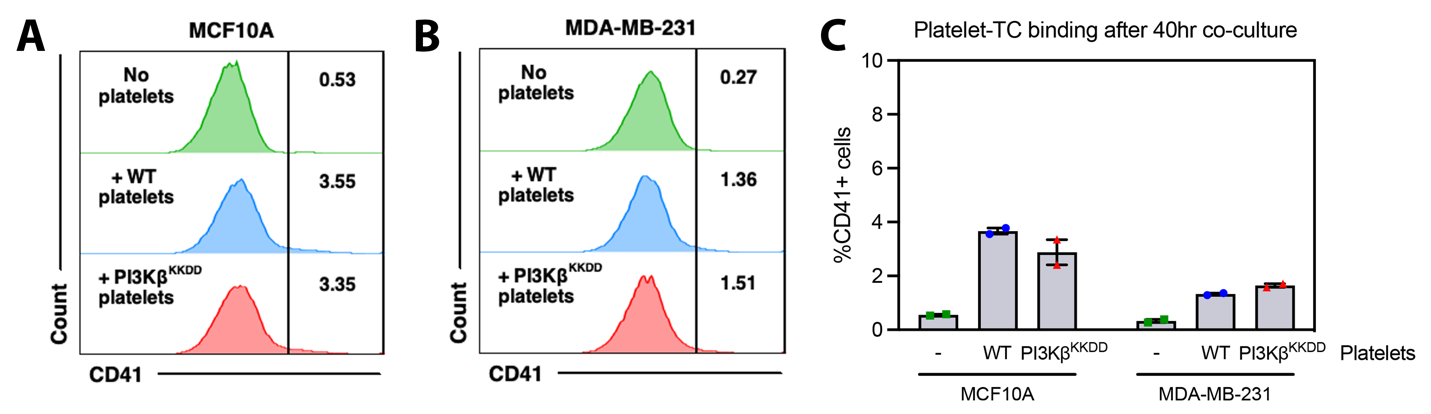

**Supplemental Figure 4. Residual platelet binding to breast epithelial or tumor cells after co-culture and washes.** MCF10A or MDA-MB-231 cells were cultured for 40 hrs without or with WT or PI3Kβ^KKDD^ platelets. After 40 hours, cells were washed three times with PBS, collected with trypsin, stained, and analyzed by flow cytometry. FACS plots show platelet interactions with MCF10A (A) or MDA-MB-231 (B) cells. (C) Quantitation of CD41+ events as a measure of platelet-tumor cell interactions. Data are presented as the mean ± SEM from n=2 independent experiments.

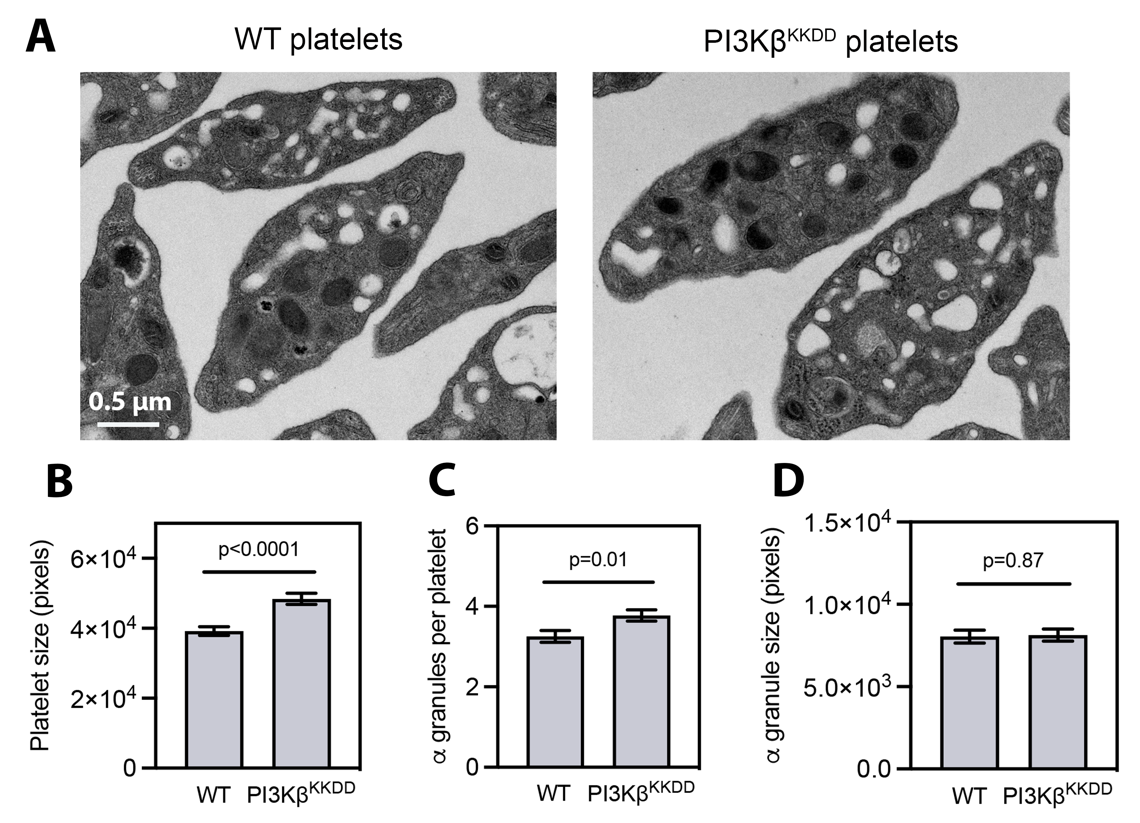

**Supplemental Figure 5. Mutation of PI3Kβ increases platelet size without affecting granule size.** (A) Representative transmission electron microscopy images of WT and PI3Kβ^KKDD^ platelets. Platelet size (B), number of alpha granules per platelet (C), and alpha granule size (D) was quantitated for each genotype. Data are presented as the mean ± SEM from at least 150 (B), 250 (C), or 50 (D) platelets from each genotype.

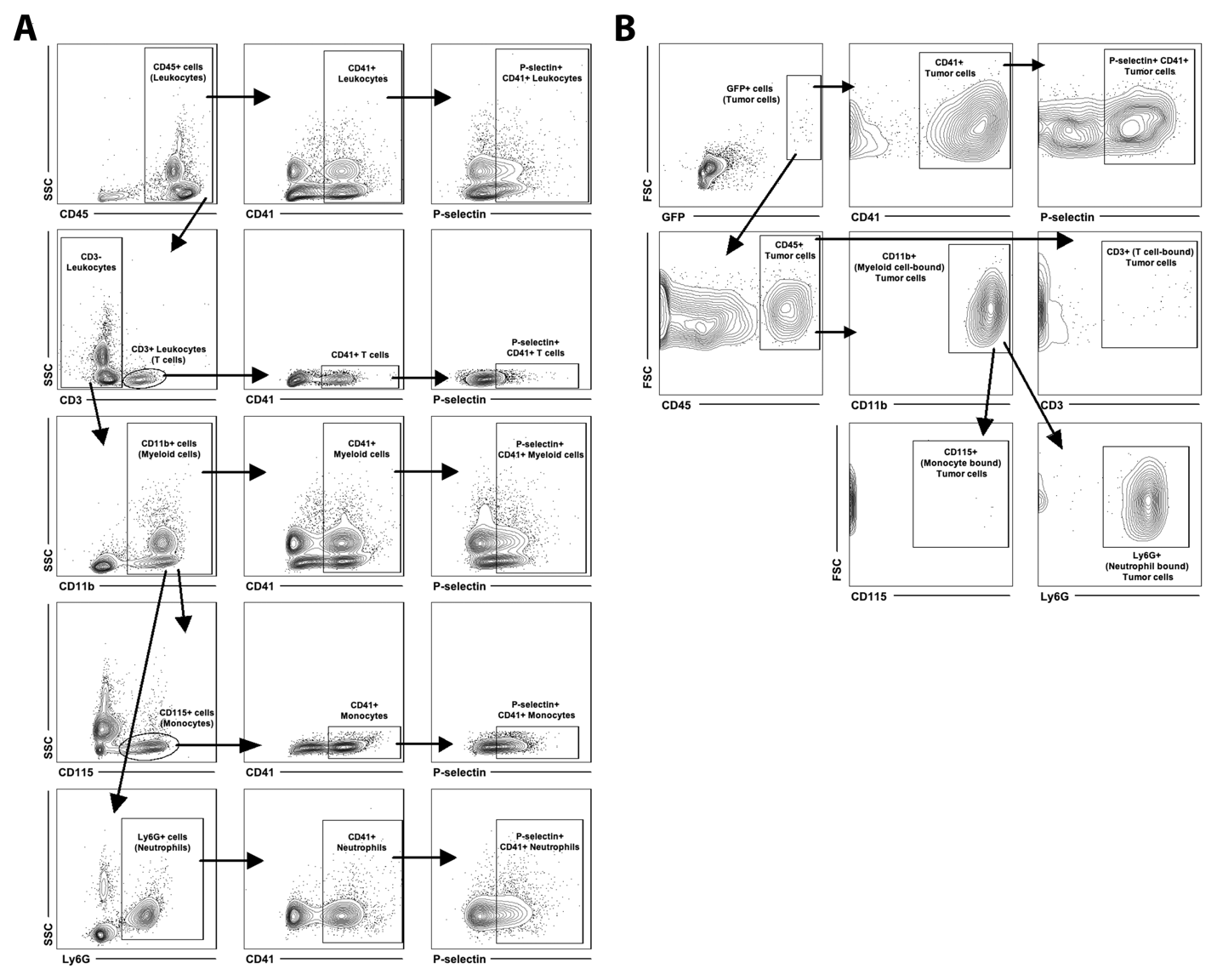

**Supplemental Figure 6. Gating scheme for flow cytometric analysis of immune cell-platelet and CTC-platelet interactions.** Blood from naïve mice or mice bearing orthotopic GFP-E0771 tumors was collected by cardiac puncture and analyzed by flow cytometry. (A) Gating strategy for defining immune cell-platelet interactions and platelet activation on immune cells. (B) Gating strategy for defining CTC-immune cell interactions, CTC-platelet interactions, and platelet activation on CTCs.

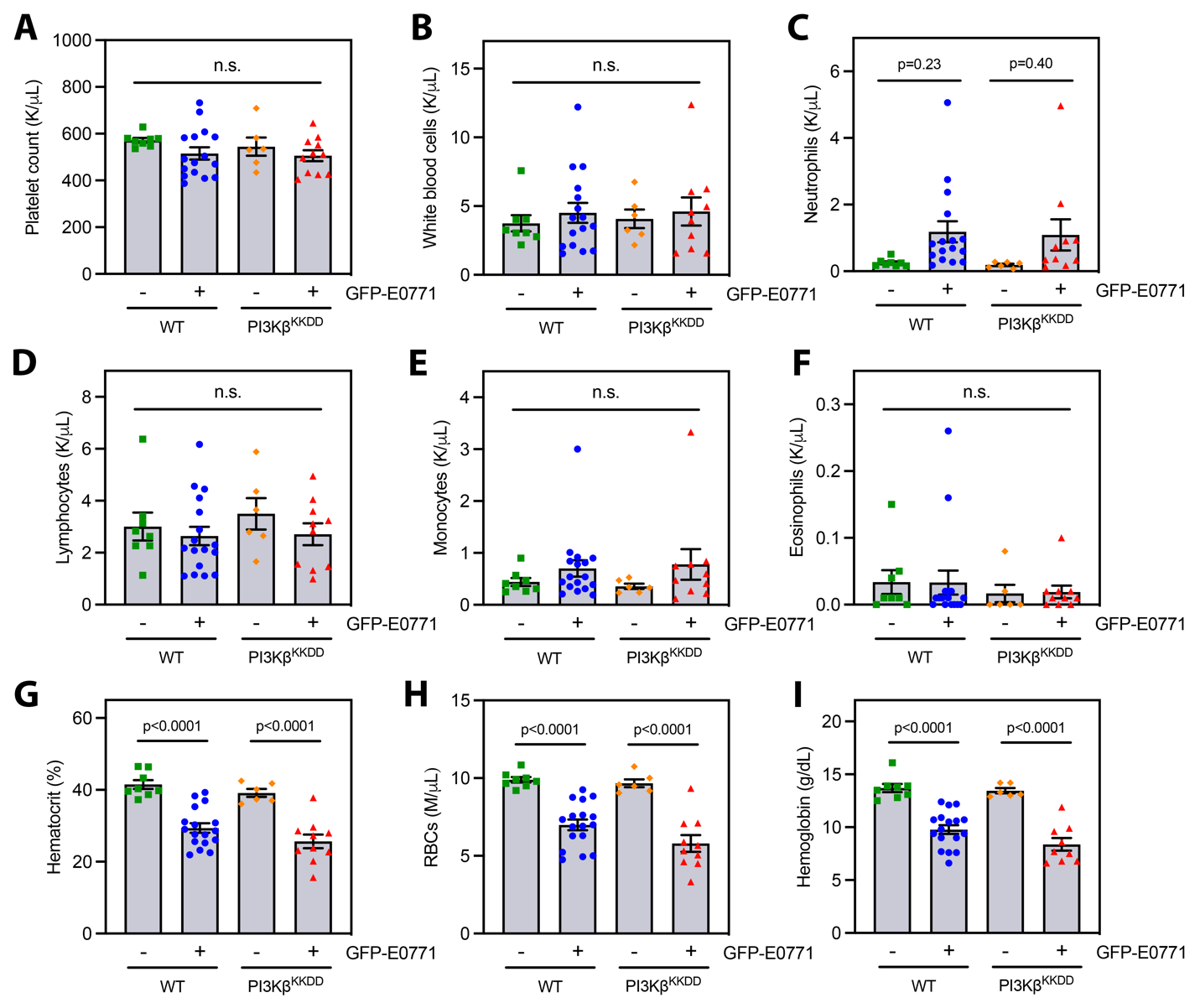

**Supplemental Figure 7. Mutation of PI3Kβ does not affect blood cell counts in naïve or tumor-bearing mice.** Blood from naïve or tumor-bearing WT or PI3Kβ^KKDD^ mice was analyzed using a hematology analyzer. The data show the numbers of platelets (A), white blood cells (B), neutrophils (C), lymphocytes (D), monocytes (E), eosinophils (F), hematocrit (G), red blood cells (H), and hemoglobin (I), and are presented as the mean ± SEM from 6-17 mice per group.
